# Host specificity in cereal rust fungi is mediated by a conserved glycoside hydrolase family

**DOI:** 10.64898/2026.05.20.726508

**Authors:** Urooj Fatima, Natalia Arango Lopez, Umar F. Shahul Hameed, Francisco J. Guzmán-Vega, Emile Cavalet-Giorsa, Naganand Rayapuram, Heribert Hirt, Michael Abrouk, Yajun Wang, Jonathan D. G. Jones, Stefan T. Arold, Simon G. Krattinger

## Abstract

Non-host resistance refers to the immunity of plant species to virtually all isolates of a potential pathogen and represents an underexplored avenue for breeding and engineering disease resistance. In domesticated and wild barley, cell surface-localized lectin receptor kinases (LecRKs) contribute to determining the host status to leaf rust fungi, which pose a major threat to global cereal production. Here, we identify a conserved family of leaf rust glycoside hydrolases as ligands for these barley LecRKs and show that direct ligand-receptor binding triggers immune responses. This mechanism of pathogen perception is conserved across multiple cereal species and can be functionally transferred between them. We also uncover previously uncharacterized recognition specificities among distinct LecRK variants, expanding the repertoire of LecRK-mediated rust pathogen detection. Our findings define a molecular mechanism underlying non-host resistance in cereals and provide a basis for harnessing non-host rust resistance across diverse crop-pathogen systems.

## Main

Rusts are destructive fungal plant diseases that affect many cereal crop species^1,2^. In wheat (*Triticum aestivum*) alone, rusts cause annual production losses of ∼48 million tons^2^. Cereal rust pathogens exhibit a high degree of host specificity^3,4^, with each rust species or *forma specialis* generally having a narrow host range and typically infecting a single cereal host. Genetic analyses in the cereal crop barley (*Hordeum vulgare*) have revealed multiple quantitative trait loci (QTL) that determine the host status of barley to various rust species^5^, leading to the identification of cell surface-localized receptor kinases^6,7^ and intracellular nucleotide-binding, leucine-rich repeat (NLR) immune receptors^8^ underlying these QTL. The orthologous lectin receptor kinases Hv-LecRK (encoded by the *Rphq2* locus^9^) from domesticated barley and Hb-LecRK (encoded by the *Rph22* locus^10^) from wild bulbous barley (*Hordeum bulbosum*) contribute quantitatively to the non-host status of barley to different leaf rust pathogens^6^. Hv-LecRK confers strong resistance to the leaf rust species adapted to wheat (*Puccinia triticina*; *Pt*) and bulbous barley (*P. hordei-bulbosi*; *Phb*) but only weak resistance to the leaf rust adapted to domesticated barley (*P. hordei*; *Ph*). Conversely, Hb-LecRK provides strong resistance to *Pt* and *Ph*, but weak resistance to *Phb* adapted to bulbous barley^6^ (Fig. 1a). Hv-LecRK and Hb-LecRK thus influence the status of barley as a host to various leaf rust species, bridging quantitative host and non-host resistance. However, their molecular function, particularly the ligands they recognize, remains unknown. Here, we identify and describe orthologous glycoside hydrolase 5 (GH5) proteins from *Ph, Phb*, and *Pt* that bind to Hv-LecRK and Hb-LecRK, activating defense responses. We further show that LecRK-mediated GH5 perception is conserved across multiple cereal species, offering the possibility of engineering non-host rust resistance through cross-species transfer of LecRKs.

**Fig. 1.**
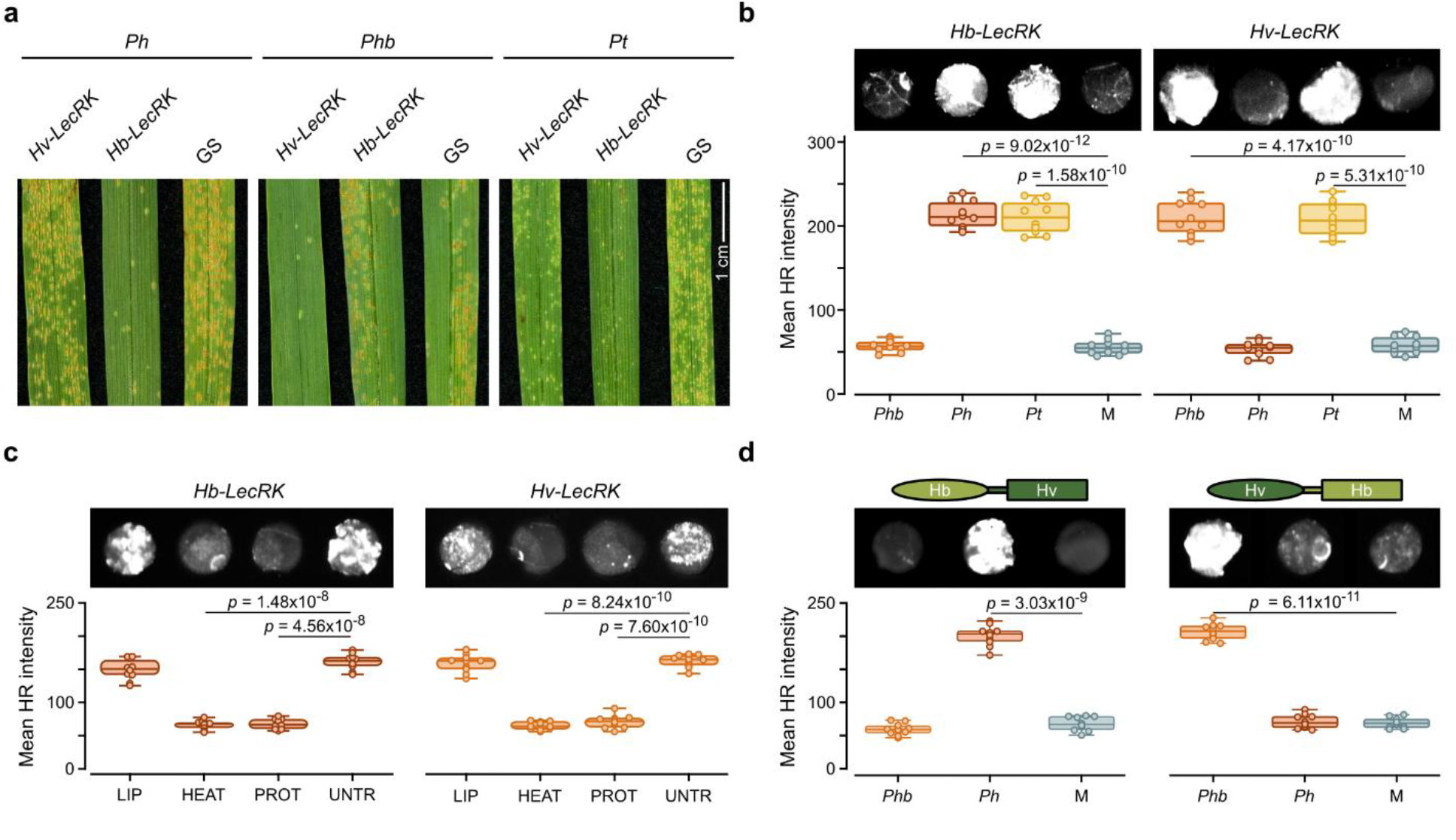
Hb-LecRK and Hv-LecRK are activated by a proteinaceous ligand from leaf rust pathogens. **a**, Representative photographs, taken 14 days post inoculation (dpi), showing the phenotypes resulting from infection by *P. hordei* (*Ph*), *P. hordei-bulbosi* (*Phb*) or *P. triticina* (*Pt*) in the leaves of transgenic barley lines expressing *Hv-LecRK* or *Hb-LecRK* in the genetic background of barley line Golden SusPtrit (GS). **b**, Quantification of the hypersensitive response (HR) induced by recognition of a ligand for LecRKs. *Hb-LecRK* or *Hv-LecRK* was expressed in leaves of *Nicotiana benthamiana* plants, which were then infiltrated with apoplastic wash fluid (AWF) from barley leaves infected with *Phb, Ph* or *Pt* and mock controls (M). **c**, HR induced by infiltration of *N. benthamiana* leaves expressing *Hb-LecRK* or *Hv-LecRK* with AWF from *Ph* or *Phb* that was pre-treated with lipase (LIP), heat or proteinases (PROT) prior to infiltration. Untreated (UNTR) AWF is shown as control. **d**, HR induced by expression of domain-swap variants between Hb-LecRK (light green; ellipse, lectin domain; rectangle, kinase domain) and Hv-LecRK (dark green) in *N. benthamiana* after infiltration with AWF from barley leaves infected with *Phb* or *Ph* or mock infected. For **b–d**, the HR-induced fluorescence was imaged (top) 2 days after AWF infiltration (Fusion-FX imager) and quantified (bottom) using ImageJ. Boxplots represent the quantification of mean HR intensity (dark orange, *Ph*; light orange, *Phb*; yellow, *Pt*) across ten biological replicates (n = 10). Circles indicate individual data points. The horizontal line denotes the median, box edges mark the first and third quartiles, and whiskers extend to the minimum and maximum values. The statistical significance of differences was determined by paired two-sample Student’s t-test (two-tailed); *P*-values are indicated above the comparisons. Protein abundance data are provided in Extended Data Fig. 1. The experiments were performed three times with consistent results each time.

## Results

### Activation of Hv-LecRK and Hb-LecRK signaling requires a leaf rust-derived protein

In this study, we wished to identify the cognate ligand(s) of the Hv-LecRK and Hb-LecRK receptors. To facilitate functional analysis, we established a transient expression assay that recapitulates the quantitative *in planta* responses mediated by Hv-LecRK and Hb-LecRK upon recognition of their ligand (Fig. 1b). Specifically, we transiently expressed *Hv-LecRK* or *Hb-LecRK* in the leaves of *Nicotiana benthamiana* plants via Agrobacterium (*Agrobacterium tumefaciens*)-mediated infiltration and then infiltrated them with apoplastic wash fluid (AWF) extracted from rust-inoculated barley leaves. The plant apoplast represents the extracellular space that serves as a critical interface for receptor kinase-mediated plant immunity. We used ‘SusPtrit’, an experimental barley line that is susceptible to non-adapted leaf rusts, including *Phb* and *Pt*, for inoculations and as a source of AWF^6,11^. AWF obtained from leaves infected with non-adapted leaf rusts triggered a strong hypersensitive response (HR) in the presence of the corresponding LecRK, which we evaluated qualitatively and quantitatively by measuring cell death-related fluorescence (Fig. 1b). In contrast, AWF obtained from barley leaves inoculated with adapted rusts (*Ph* for Hv-LecRK; *Phb* for Hb-LecRK) or from mock-inoculated leaves failed to induce HR (Fig. 1b). Pre-treating AWF with heat or proteinases abolished HR (Fig. 1c), indicating that the HR depends on a proteinaceous leaf rust-derived factor. Additionally, AWF extracted from barley leaves inoculated with wheat stripe rust (*P. striiformis* f. sp. *tritici*; *Pst*) or wheat stem rust (*P. graminis* f. sp. *tritici*; *Pgt*) did not trigger HR (Extended Data Fig. 1), consistent with the specificity of *Rphq2* and *Rph22* to leaf rusts^6^. A domain-swap experiment between Hv-LecRK and Hb-LecRK revealed that the extracellular lectin domains, not the kinase domains, govern the strength of HR against different leaf rust pathogens (Fig. 1d). These results indicate that the resistance mechanism of Hv-LecRK and Hb-LecRK involves a leaf rust-derived protein, possibly acting as ligand.

### Conserved leaf rust glycoside hydrolases activate Hv-LecRK and Hb-LecRK-mediated defense responses

To identify apoplastic candidate leaf rust proteins, we performed liquid chromatography-mass spectrometry (LC-MS) on AWF extracted from SusPtrit leaves inoculated with *Ph* or *Phb*. We collected AWF from inoculated and mock-treated leaves at 4 days post inoculation (dpi). We then assigned detected peptides to predicted proteins from leaf rust or barley using genome annotations from the assemblies of *Ph* isolate *Ph560*^12^ and the barley cultivar ‘Vada’ (Supplementary Note 1). After quality filtering, we retained 232 predicted *Ph* proteins (Supplementary Table 1). To prioritize candidate ligands for LecRK recognition, we used AlphaFold2^13^ to predict putative complexes between each of the 232 *Ph* proteins and Hv-LecRK or Hb-LecRK, ranking interactions by their ipTM scores (Supplementary Tables 1 and 2). Based on these predictions, we synthesized constructs encoding the top 37 candidates based on the *Ph560* genome sequence and transiently co-expressed them in *N. benthamiana* leaves along with *Hv-LecRK* or *Hb-LecRK*. Among them, one construct, encoding a predicted leaf rust glycoside hydrolase of 519 amino acids from the GH5 family with predicted endo-β-1,6-glucosidase activity, triggered HR exclusively in the presence of Hb-LecRK (Extended Data Fig. 2). One *Ph* construct induced an HR even without Hb-LecRK, indicating a possible detection by an endogenous *Nicotiana* receptor, while the remaining 35 *Ph* constructs failed to elicit HR when co-infiltrated with the *Hb-LecRK* gene (Extended Data Fig. 2). The putative leaf rust GH5 protein ranked fourth and thirteenth among the 232 predicted *Ph* apoplastic proteins for interaction with Hv-LecRK and Hb-LecRK, respectively, based on AlphaFold2 ipTM scores (Supplementary Tables 1 and 2).

We next obtained the sequences of the GH5-encoding gene from *Ph* isolate 1.2.1 (the isolate used for experiments in this study), as well as its orthologs from *Phb* (*GH5*^*Phb*^), *Pt* (*GH5*^*Pt*^), *Pst* (*GH5*^*Pst*^) and *Pgt* (*GH5*^*Pgt*^), through genome sequencing and searching of public databases. *GH5*^*Ph*^ from *Ph* isolate 1.2.1 was identical to the version from the *Ph560* reference assembly. The *Phb* genome carried two genes orthologous to *GH5*^*Ph*^, named *GH5*^*Phb_h1*^ and *GH5*^*Phb_h2*^, possibly representing haplotypes of the two separated rust nuclei. GH5^Ph^ differed by five amino acids and three amino acids from GH5^Phb_h1^ and GH5^Phb_h2^, respectively (Fig. 2a). We validated the predicted coding sequences of *GH5*^*Ph*^, *GH5*^*Phb_h1*^ and *GH5*^*Phb_h2*^ using full-length transcriptome sequencing (Iso-Seq), confirming that the predicted GH5 protein variants are all 519 amino acids in length. *GH5*^*Ph*^ expression induced a strong HR in *N. benthamiana* leaves in the presence of Hb-LecRK, but not Hv-LecRK (Fig. 2b). Conversely, *GH5*^*Phb_h1*^ expression induced HR only in the presence of Hv-LecRK, but not Hb-LecRK. An amino acid swap experiment between GH5^Ph^ and GH5^Phb_h1^ point to the polymorphic amino acid residues 60 and 458 as contributing to recognition specificity (Extended Data Fig. 3). *GH5*^*Pt*^ expression from wheat leaf rust activated a strong HR with both receptors, mirroring the rust phenotype observed in barley^6^ (Figs. 1a and 2b). The GH5 versions from *Pst* and *Pgt* did not activate HR with either receptor kinase, in agreement with the fact that *Rphq2* and *Rph22* do not confer resistance to diseases other than leaf rust (Extended Data Fig. 4). GH5^Phb_h2^, the second GH5 variant present in *Phb*, showed a response similar to GH5^Ph^, activating a strong HR in the presence of Hb-LecRK, but not Hv-LecRK (Fig. 2b).

**Fig. 2.**
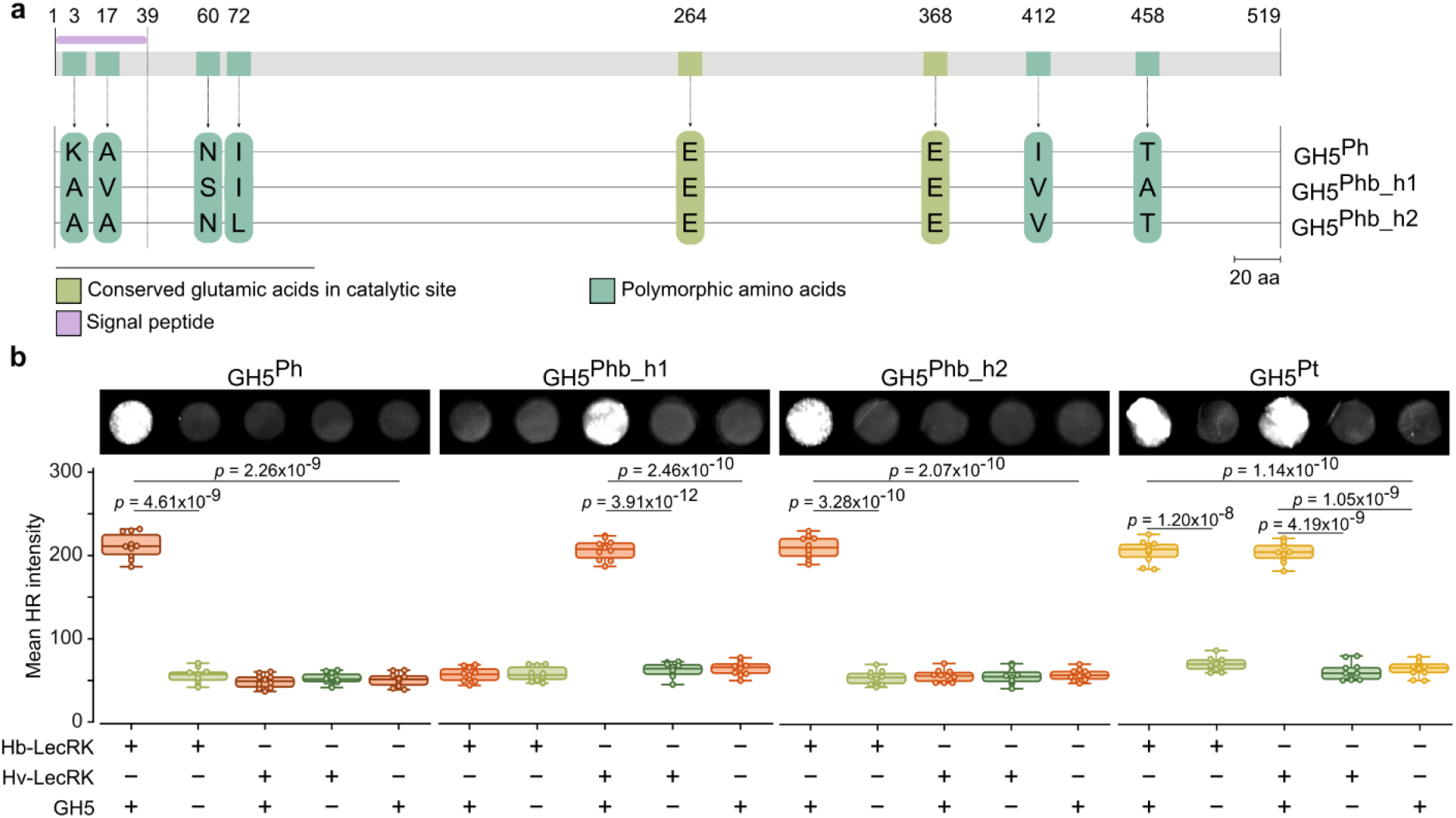
Differential recognition of leaf rust GH5 variants by Hv-LecRK and Hb-LecRK mirrors pathogen specificity. **a**, Diagram of the GH5 variants from *Ph* and *Phb*, with the amino acid differences across variants and the conserved amino acids in the catalytic site shown. Numbers indicate the amino acid numbers. **b**, HR imaging (top) and quantification (bottom) in *N. benthamiana* leaves co-expressing *GH5* variants from different rust species and *Hb-LecRK* or *Hv-LecRK*. (+) indicates co-infiltration of the *LecRK* construct with the *GH5* construct; (-) indicates infiltration of the *LecRK* construct alone; ‘No LecRK’ indicates GH5 infiltrated alone. HR was imaged 3 dpi (Fusion-FX imager). Box plots represent the quantification of the mean HR intensity of ten biological replicates (*n* = 10). Circles indicate individual data points. The horizontal line denotes the median, box edges mark the first and third quartiles, and whiskers extend to the minimum and maximum values. The statistical significance of differences was determined by paired two-sample Student’s *t*-test (two-tailed); *P*-values are indicated above the comparisons. Protein abundance data are provided in Extended Data Fig. 4b. The experiments were performed three times with consistent results.

The identified GH5 proteins are predicted to possess endo-β-1,6-glucosidase activity. β-1,6-glucan is a component specifically found in fungal cell walls^14^. The endogenous function of the GH5 members identified in this study might therefore involve the modification of fungal cell wall during invasive growth or the degradation of fungal cell-wall components that may serve as pathogen-associated molecular patterns (PAMPs)^14,15^. Enzyme activity assays confirmed β-1,6-glucosidase activity for GH5^Ph^, GH5^Phb_h1^ and GH5^Phb_h2^ (Extended Data Fig. 5a, b). GH5 proteins contain two conserved glutamic acid (E) residues that are predicted to be critical for catalytic activity^16,17^. A GH5^Ph^ variant carrying mutations in these two predicted catalytic residues lost β-1,6-glucosidase activity (Extended Data Fig. 5a, b) but still elicited HR in the presence of Hb-LecRK (Extended Data Fig. 5c), indicating that recognition by the lectin receptor kinase is independent of the GH5 catalytic activity of its putative ligand.

### Leaf rust GH5 proteins directly bind to Hv-LecRK and Hb-LecRK

To test interactions between leaf rust GH5 proteins and Hv-LecRK or Hb-LecRK *in planta*, we performed co-immunoprecipitation (co-IP) assays in *N. benthamiana* leaves co-expressing each full-length *LecRK* and *GH5* variants. Myc-tagged GH5^Ph^ and Myc-tagged GH5^Phb_h2^ co-precipitated more strongly with full-length Hb-LecRK tagged with the green fluorescent protein (Hb-LecRK-GFP) than did Myc-tagged GH5^Phb_h1^, whereas Hv-LecRK-GFP showed a stronger interaction with Myc-tagged GH5^Phb_h1^ than with Myc-tagged GH5^Ph^ or Myc-tagged GH5^Phb_h2^ following immunoprecipitation with anti-GFP beads (Fig. 3a). HiBiT-based co-IP assays further quantified relative interaction strengths and confirmed the interaction specificity between the GH5 variants and the two LecRKs (Fig. 3b, Extended Data Fig. 6a, b). In contrast, Myc-tagged GH5^Pt^ exhibited strong interactions with both Hv-LecRK-GFP and Hb-LecRK-GFP as quantified by HiBiT-based co-IP (Extended Data Fig. 6c-e). Furthermore, *in planta* co-IP with a Myc-tagged catalytically inactive GH5^Ph^ mutants retained the same interaction specificity as wild-type Myc-tagged GH5^Ph^, indicating that LecRK-mediated perception is independent of the GH5 catalytic activity (Extended Data Fig. 7a).

**Fig. 3.**
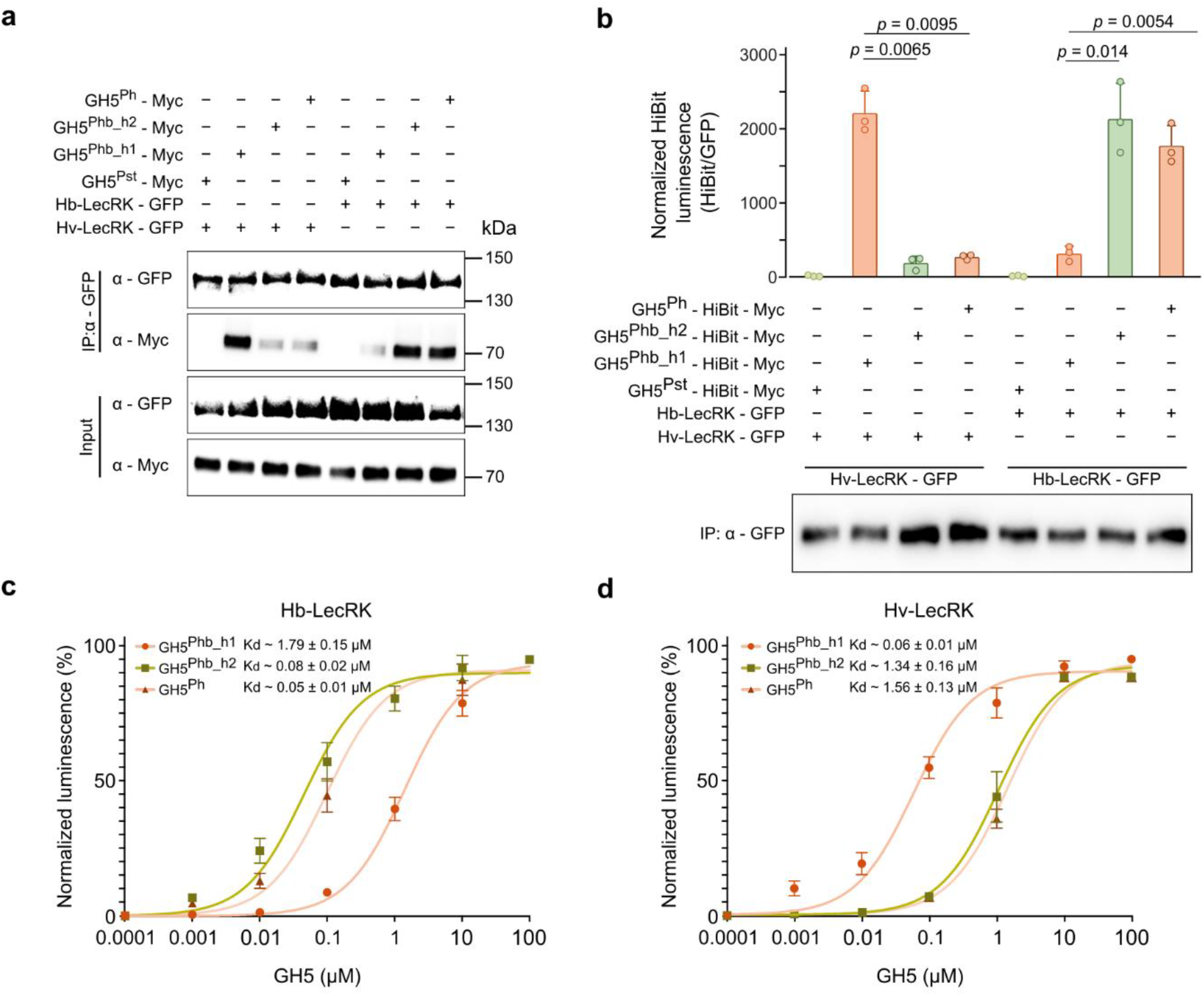
Direct binding of GH5^Ph^, GH5^Phb_h1^ and GH5^Phb_h2^ to Hv-LecRK and Hb-LecRK. **a**, *In planta* co-immunoprecipitation (co-IP) assay of Myc-tagged GH5^Ph^, GH5^Phb_h2^, GH5^Phb_h1^ and GH5^Pst^ (as negative control) transiently co-expressed in *N. benthamiana* leaves with GFP-tagged full-length Hv-LecRK or Hb-LecRK. Total protein extracts were immunoprecipitated with anti-GFP beads (IP) and probed with anti-Myc and anti-GFP antibodies. **b**, HiBiT-based quantitative co-IP assay of Myc-HiBiT-tagged GH5^Ph^, GH5^Phb_h2^, GH5^Phb_h1^ and GH5^Pst^ (as negative control) transiently co-expressed in *N. benthamiana* leaves with GFP-tagged full-length Hv-LecRK or Hb-LecRK. Total protein extracts were subjected to immunoprecipitation (IP) using anti-GFP beads. HiBiT luminescence (**b**, top) was measured from IP samples and normalized to the abundance of immunoprecipitated Hv-LecRK or Hb-LecRK, quantified by ImageJ analysis of anti-GFP immunoblots of the corresponding IP samples (**b**, bottom). Data are presented as mean ± s.d. normalized luminescence intensities from three independent experiments (n = 3). Circles indicate individual data points. The statistical significance of differences was determined by paired two-sample Student’s *t*-test (two-tailed); *P*-values are indicated above the comparisons. Absolute luminescence values after background subtraction for IP and input samples, together with protein abundance detected by immunoblotting with anti-Myc and anti-GFP antibodies, are shown in Extended Data Fig. 6a, b. **c–d**, Binding assay of GFP-tagged Hb-LecRK (**c**), or Hv-LecRK (**d**) with HiBit-tagged GH5^Ph^, GH5^Phb_h1^ and GH5^Phb_h2^ proteins purified from the apoplastic wash fluid of *N. benthamiana* leaves. Binding was quantified by measuring HiBiT luminescence following incubation of receptor proteins with increasing concentrations of GH5 proteins. Data are presented as the percentage of maximal HiBiT luminescence obtained at saturating concentrations of each GH5 protein. Values represent mean ± s.e.m. from three independent experiments, with four technical replicates each. Binding curves were fitted to a rectangular hyperbola model to determine apparent dissociation constant (*K*_*d*_) values. GFP incubated with GH5^Ph^, GH5^Phb_h1^ and GH5^Phb_h2^ was used as a negative control and the corresponding binding curves are shown in Extended Data Fig. 6f. The experiments were performed three times with consistent results.

To determine whether the interactions detected by co-IP reflect direct binding between GH5 proteins and LecRKs, we performed quantitative binding assays using the NanoLuc-based HiBiT–LgBiT complementation system with proteins purified from *N. benthamiana*^18,19^. Purified HiBiT-tagged GH5 proteins from *N. benthamiana* AWF were incubated with immobilized GFP-tagged LecRKs purified from *N. benthamiana* leaves, and binding affinities were determined by saturation binding analysis using increasing concentrations of GH5 proteins. These assays demonstrated direct and selective binding of GH5 variants to the two LecRKs (Fig. 3c, d). GH5^Ph^ and GH5^Phb_h2^ bound Hb-LecRK with substantially higher affinity, with apparent dissociation constants (*K*_d_) of 0.050 ± 0.01 µM and 0.060 ± 0.02 µM, respectively than GH5^Phb_h1^ (*K*_d_ = 1.79 ± 0.15 µM) (Fig. 3c). Conversely, Hv-LecRK displayed high affinity for GH5^Phb_h1^ (*K*_d_ = 0.06 ± 0.01 µM), whereas binding to GH5^Ph^ and GH5^Phb_h2^ was markedly weaker (*K*_d_ = 1.56 ± 0.13 µM and 1.34 ± 0.16 µM, respectively) (Fig. 3d). These quantitative differences in binding affinity closely paralleled the differential resistance phenotypes observed *in planta*, supporting that recognition by Hv-LecRK and Hb-LecRK is determined by preferential GH5 binding.

Together, these results indicate a direct interaction between the barley lectin receptor kinases and the leaf rust-derived GH5 variants. The results obtained with GH5^Phb_h2^ are noteworthy: to our knowledge, only two isolates of *Phb* have been reported in the literature^20^. Compared to *Ph* on barley, *Phb* grows relatively slowly on *H. bulbosum*. This observation raises the possibility that *Phb* may have additional hosts, reflecting a tendency for pathogens that infect wild crop relatives in natural ecosystems to be generally less virulent and more generalist than the highly virulent and specialized pathogens that infect domesticated crops in agricultural ecosystems^21^.

### A rice blast pathogen expressing leaf rust *GH5* variants activates resistance in barley expressing *Hv-LecRK* and *Hb-LecRK*

Because rust pathogens are recalcitrant to transformation, we transformed the sequence encoding the GH5 variants from *Ph* and *Phb* into the rice blast-conferring pathogen *Magnaporthe oryzae*, which can also infect barley. *M. oryzae* transformants expressing *GH5*^*Ph*^ triggered resistance on *Hb-LecRK*-expressing transgenic barley lines, but not on lines with *Hv-LecRK* or the non-transgenic wild-type (Fig. 4a). Conversely, *M. oryzae* expressing *GH5*^*Phb_h1*^ resulted in resistance only on *Hv-LecRK*-expressing barley lines (Fig. 4b), while *M. oryzae* strains expressing *GH5*^*Phb_h2*^ induced resistance responses comparable to those triggered by *M. oryzae* expressing *GH5*^*Ph*^ (Fig. 4c). All barley lines were susceptible to *M. oryzae* strains carrying the empty vector (Fig. 4d). *M. oryzae* transformants expressing the catalytically inactive *GH5*^*Ph*^ variant (*m3GH5*^*Ph*^) triggered resistance on *Hb-LecRK*-expressing transgenic barley lines, but not on *Hv-LecRK* lines or the non-transgenic plants, similar to transformants expressing wild-type *GH5*^*Ph*^ (Extended Data Fig. 7b), confirming that LecRK-mediated resistance is independent of GH5^Ph^ catalytic activity.

**Fig. 4.**
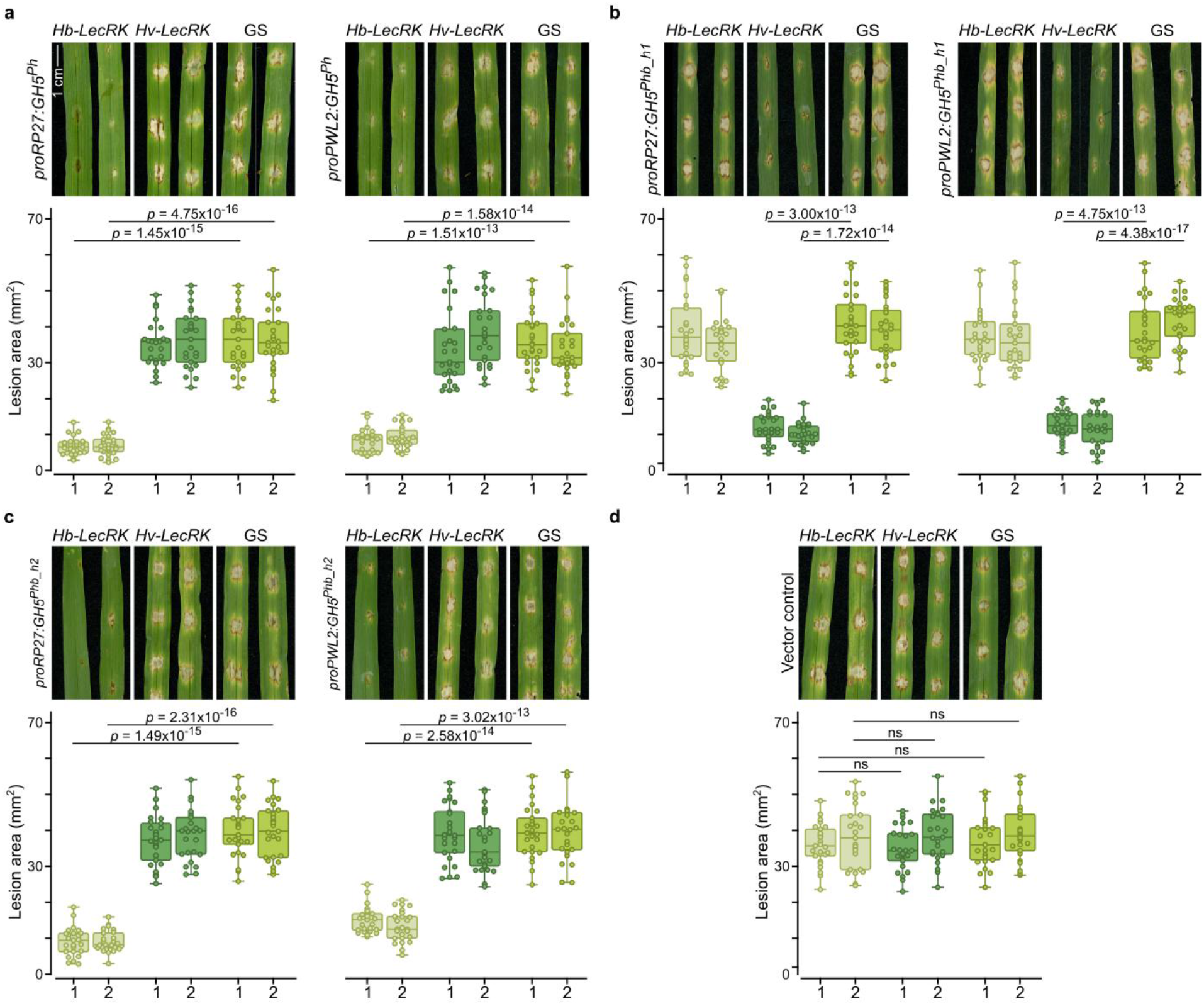
*GH5* variants expressed in *M. oryzae* reveal Hv-LecRK and Hb-LecRK specificity in barley. **a–d**, Representative photographs of (top) and lesion areas from (bottom) the leaves of barley transgenic lines expressing *Hb-LecRK* or *Hv-LecRK*, or the wild type Golden SusPtrit (GS), after inoculation with *M. oryzae* expressing *GH5*^*Ph*^ (**a**), *GH5*^*Phb_h1*^ (**b**) or *GH5*^*Phb_h2*^ (**c**) driven by the *RP27* (left) or *PWL2* (right) promoter, or with *M. oryzae* carrying the empty vector (**d**). Leaves were inoculated using the spot inoculation method. Numbers 1 and 2 on the *x*-axis represent two independent fungal transformants. Spot inoculations were performed on intact barley leaves without pricking, and photographs were taken at 6 dpi. Boxplots represent the mean lesion area (*n* = 25) obtained from five leaves collected from five independent biological replicates for each treatment and experiment. Circles denote individual data points. The horizontal line denotes the median, box edges mark the first and third quartiles, and whiskers extend to the minimum and maximum values. The statistical significance of differences was determined by paired two-sample Student’s *t*-test (two-tailed); *P*-values are indicated above the comparisons. ns, not significantly different. The experiments were performed three times with consistent results.

### LecRK-mediated resistance shows cross-species conservation and transferability

Phylogenetic analysis across seven rust species revealed that GH5^Ph^, GH5^Phb_h1^ and GH5^Phb_h2^ are members of a conserved clade (Fig. 5a), raising the possibility that LecRK – GH5 interactions, mediating non-host resistance, may be more broadly conserved.

**Fig. 5.**
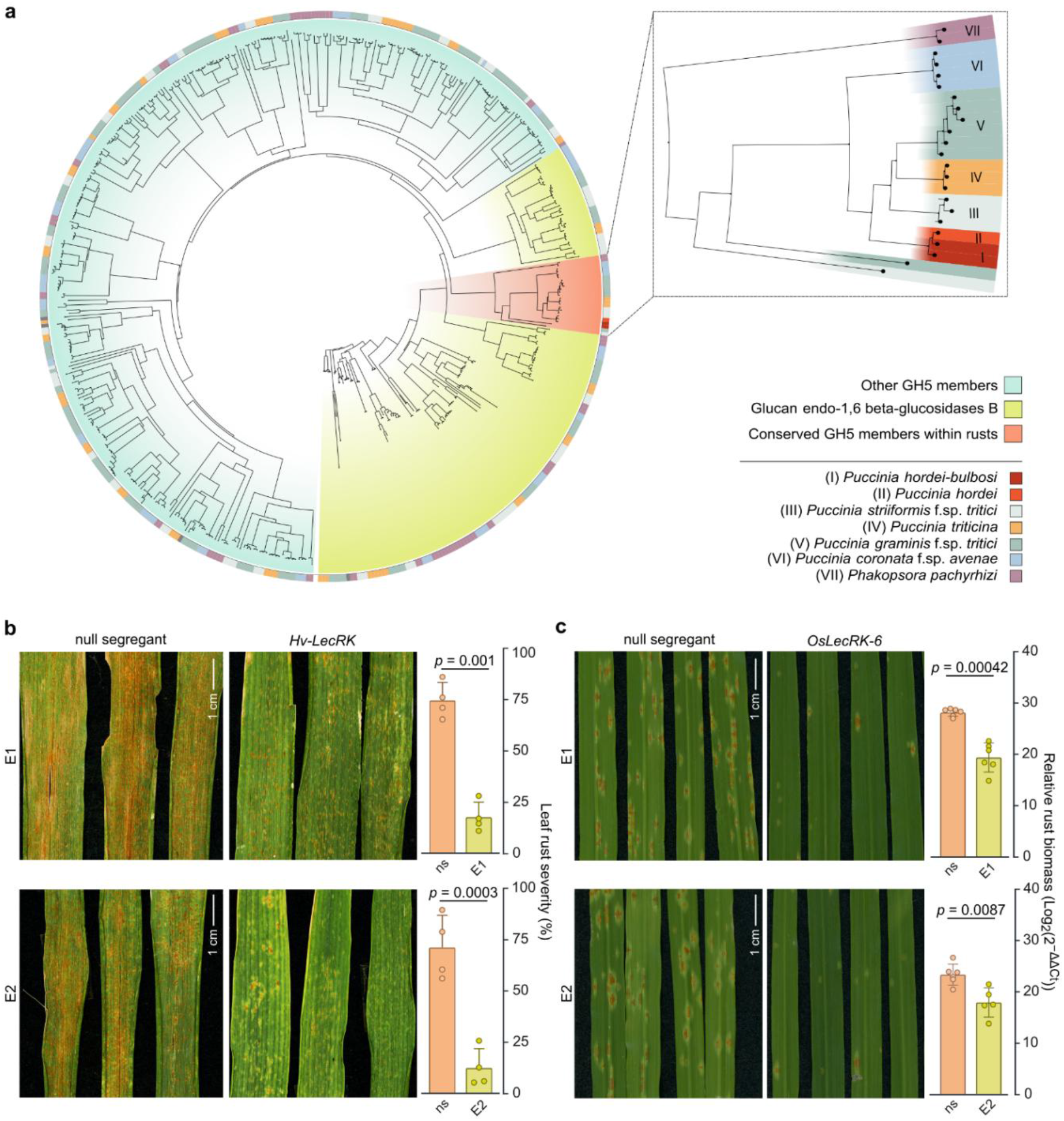
Conservation and transferability of GH5-mediated rust perception across cereal species. **a**, Phylogenetic analysis of GH5 members from seven different rust species. All protein sequences were extracted from the Carbohydrate Active Enzymes database (CAZy). **b**, Representative photographs (left) and quantification (right) of leaf rust severity on the flag leaves from wheat transgenic lines expressing *Hv-LecRK* and corresponding null segregants (ns) grown on the KAUST experimental field site. Disease severity was assessed 70 days after inoculation of the spreader rows with a mixture of seven *Pt* isolates. Values represent mean ± s.d. from four independent rows, each containing 15 plants (n = 60). The statistical significance of differences was determined by paired two-sample Student’s *t*-test (two-tailed). E1 (top) and E2 (bottom) represent two independent transgenic events. Field studies were conducted for two consecutive years; the data presented here is from year one. **c**, Left, representative photographs of leaves from adult wheat transgenic lines expressing *OsLecRK-6* or their null segregants after infection with *Pgt*. The photographs were taken at 11 dpi for six independent biological replicates (*n =* 6) and two independent events (E1 and E2). Inoculations were done in controlled conditions. Right, relative fungal biomass quantified in *OsLecRK-6* transgenic lines and null segregants using qPCR (*n =* 5–6). Values are shown as means ± standard deviation. The statistical significance of differences was determined by paired two-sample Student’s *t*-test (two-tailed). The experiments were performed three times with consistent results each time.

Introducing *Hv-LecRK* or *Hb-LecRK* into wheat cultivar Fielder increased resistance against a mixture of seven *Pt* isolates in field trials conducted across two seasons (Fig. 5b, Extended Data Figs. 8 and 9), demonstrating that LecRK-mediated rust resistance is transferable across species.

However, Hv-LecRK and Hb-LecRK perceived only leaf rust-derived GH5 variants, but not orthologous variants from other rust species. To assess the extent of conservation of the LecRK-mediated GH5 perception and the possibility of an expanded GH5 recognition spectrum, we screened multiple combinations of rice (*Oryza sativa*) LecRKs and GH5 orthologs from various rust species (Supplementary Table 3) using the *N. benthamiana* system (Extended Data Fig. 10a). In particular, we focused on the rice cultivar ‘Nipponbare’, which carries a LecRK cluster containing ten paralogous copies, whereas Hv-LecRK is a single-copy gene in barley. Several rice LecRKs triggered HR when their encoding constructs were co-infiltrated with a *GH5*^*Pgt*^ construct from wheat stem rust, suggesting that they have an expanded recognition spectrum (Extended Data Fig. 10a, b). We introduced a construct encoding one of these rice LecRKs, *OsLecRK-6* (LOC_Os08g03090.1), into wheat cultivar Fielder. The wheat transgenic lines expressing *OsLecRK-6* were more resistant to *Pgt* than their respective null-segregants (Fig. 5c), which was supported by a significantly lower relative fungal biomass accumulating in these *OsLecRK-6-* expressing transgenic lines (Fig. 5c). Co-IP assays indicated an interaction between GH5^Pgt^ and OsLecRK-6, but not between GH5^Pgt^ and Hv-LecRK or Hb-LecRK (Extended Data Fig. 10c). These results suggest that GH5 recognition by LecRKs represents a molecular resistance mechanism conserved across multiple cereal crop species that can be harnessed for disease resistance breeding and engineering.

## Discussion

Non-host resistance is sometimes described as durable and broad-spectrum^3,4,22,23,24^ because it ideally acts against all genetic variants of a potential pathogen. Despite its potential importance for disease resistance breeding and agriculture, the genetic and molecular underpinnings of non-host resistance remain poorly understood, largely due to the challenges of genetically dissecting non-host resistance for different plant-pathosystems. In fact, the molecular basis of non-host resistance has been identified as one of the top ten unanswered questions in molecular plant-microbe interactions^25^. The cereal rust - barley pathosystem, characterized by a high degree of host-specialization and the near non-host status of barley to different rust pathogens, provides an excellent model to address this question and unravel the genetic and molecular mechanisms that can result in non-host resistance. Studies in barley and other plant species have shown that, similar to host resistance, non-host resistance is mediated by both cell surface-localized pattern recognition receptors (PRRs) and intracellular NLR immune receptors^6-8,26^. This overlap in molecular components of host and non-host resistance raises a central question: Are genes conferring non-host resistance inherently more durable than those mediating host resistance?

Here, we significantly advanced the molecular understanding of non-host resistance against cereal rusts. We show that leaf rusts produce proteins belonging to glycoside hydrolases of the GH5 family and that these proteins are highly conserved across different rust species and perceived by LecRKs. The high conservation of the GH5 proteins perceived by Hv-LecRK and Hb-LecRK across different leaf rust species and other rust taxa suggests that this GH5 clade may play a role during infection progression, potentially resulting in evolutionary constraints that may limit diversification and thus diminish the likelihood of a rapid breakdown of *Rphq2*- and *Rph22*-mediated resistance. Nonetheless, the weaker resistance observed with adapted leaf rust species (*Ph* with Hv-LecRK and *Phb* with Hb-LecRK) indicates that some level of evasion has evolved, possibly through reduced GH5-LecRK binding affinity (Fig. 3c, d). In addition, there appears to be functional variation among *LecRK* alleles in barley. SusPtrit, which is susceptible to multiple rust species, carries a *LecRK* allele (*Hv*-*LecRK*^*SusPtrit*^) encoding a protein that does not recognize GH5^Ph^, GH5^Phb^ or GH5^Pt^ (Extended Data Fig. 10d). Knowing when LecRK-GH5 interactions are functional could help deploy functional LecRK versions in breeding, reducing the risk of host jumps. For example, the host jump of the blast fungus to wheat was facilitated by the widespread deployment of wheat cultivars that lacked a key blast resistance gene involved in determining host specificity^27^. Together, these molecular insights into the LecRK-GH5 interaction, and the conservation of this recognition mechanism across different cereal species, provide a foundation for transferring or engineering LecRK variants with broadened or altered recognition specificities across multiple crops.

## Acknowledgements

This research used the Ibex high performance cluster managed by the Supercomputing Core Laboratory at King Abdullah University of Science and Technology (KAUST). We thank the systems administrators and computational scientists for help with debugging and overall support. We are grateful to Elisabet Poquet Faig and Xingfang Shi (KAUST) for greenhouse assistance, Kymbat Zhakupova (KAUST) for wheat transformations, Kit Xi Liew for assistance with LC-MS sample runs and analyses, and Jack Rhodes (The Sainsbury Laboatory) for feedback on the manuscript. This publication is based upon work supported by KAUST awards OSR-CRG2019-4038 to U.F., N.R., H.H. and S.G.K. and OSR-OFP2023-1229 to U.F., N.A.L. and S.G.K.

## Author contributions

U.F., S.T.A., J.D.G.J. and S.G.K. conceived the project and designed the study. U.F. established the *N. benthamiana* transient assay and HR quantification, performed co-IP assays, leaf rust pathogen assays in barley and wheat transgenic lines in controlled and field conditions, and *M. oryzae* transformations and pathogen assays in barley transgenic lines. N.A. and U.F. performed transient assays for rice LecRK and stem rust pathogen assays in wheat transgenic lines. N.A. performed the phylogenetic analysis of GH5s. U.F. and U.S.H. performed the binding assays. U.F. and N.R. performed LC-MS and data analyses. F.J.G. and S.T.A. performed AlphaFold modelling. E.C.G. and U.F. performed *Ph* and *Phb* genome sequencing. M.A. and Y.W. assembled the genome sequences of the barley cultivar ‘Vada’. U.F., N.A., E.C.G. and S.K.G. designed figures. U.F. and S.G.K. drafted the first version of manuscript with input from N.A., U.F., J.D.G.J. and S.T.A. All authors read, commented on and approved the final version of the manuscript.

## Methods

### Plant material and growth conditions

Barley lines used in this study included SusPtrit, Golden SusPtrit, and the transgenic *Hv-LecRK-* and *Hb-LecRK-*expressing lines in the Golden SusPtrit background, developed in our previous research^6^. Wheat lines included cultivar Fielder and the *Hv-LecRK-* and *OsLecRK-6*-expressing transgenic lines in the Fielder background. Barley and wheat grains were sown in 9 × 9 × 9 cm pots arranged within 55 × 28 cm trays and maintained in a growth cabinet at 20 °C under a 16-hour light/8-hour dark photoperiod. *Nicotiana benthamiana* plants were grown in a growth room at 23 °C with the same 16-h light/8-h dark cycle.

### Pathogen inoculations

The rust pathogen species/isolates used in this study were *Puccinia hordei* (*Ph*) 1.2.1, a monospore derivative of isolate 1.2^28^, *P. hordei-bulbosi* (*Phb*), *P. triticina* (*Pt*) INRA^6, 29^, *P. striiformi*s f. sp. *tritici* (*Pst*), and *P. graminis* f. sp. *tritici* (*Pgt*) ^30^. Detailed information on the isolates is provided in Supplementary Table 4. Each isolate was propagated on its respective host (*Ph* on *Hordeum vulgare, Phb* on *H. bulbosum*, and *Pt, Pst, Pgt* on *Triticum aestivum*) before inoculation on the highly susceptible barley line SusPtrit.

For apoplastic wash fluid extraction, three to four SusPtrit grains were sown in 9 × 9 × 9 cm pots within 54 × 27 × 6 cm trays. Rust inoculations were performed 12 days after sowing. A mix of 50 mg of uredinospores in 20 ml of Novec™ 7100 oil was air-sprayed onto the plants inside a rust inoculation cabinet. Plants were subsequently incubated overnight in a dew chamber at 18 °C in the dark for spore germination and then transferred to a growth room at 20 °C with a 16-h light/8-h dark cycle until fleck symptoms appeared^6^. For phenotyping, adult *Hb-LecRK* and *Hv-LecRK* transgenic barley and wheat lines were inoculated with 5 mg uredinospores in 10 ml Novec™ 7100, incubated overnight in a dew chamber, and then returned to the growth room. Disease symptoms were assessed 14 days post-inoculation by scanning leaves using an Epson Perfection V600 Photo scanner. The *Pgt* inoculations were performed on adult wheat transgenic plants at the flag leaf stage using the same spray method, and the penultimate leaf was scanned for recording disease symptoms 11 days post-inoculation.

*Magnaporthe oryzae* wild-type strains (FR13) ^31^ were stored on dried filter papers at −20 °C and cultured on solid complete medium (CM) agar plates at 24 °C for up to 10 days under a 12 h photoperiod in a growth cabinet^32^. Conidia were harvested and conidial suspensions were adjusted to 5 × 10^4^ conidia/ml in a solution containing 0.2% (v/v) gelatin. For drop inoculations, barley grains were sown in two rows on both sides of 55 × 28 cm trays and maintained in a growth cabinet at 20 °C. Ten days after sowing, leaves were pinned to the soil with the adaxial side facing upward, and 20 µl droplets of conidial suspension were placed on each leaf at different positions without pricking. Plants were incubated 24 h in a dew chamber, then returned to a growth chamber at 24 °C under a 16-h light/8-h dark photoperiod for 5 days. Mock inoculations were performed using 0.02% (v/v) gelatin. Disease symptoms were recorded 6 days post-inoculation by scanning the leaves.

### Genome and transcriptome sequencing

SusPtrit grains were sown (4-5 per pot, 6 × 6 × 10 cm) in a growth chamber at 20 °C under long-day photoperiods of 16-h day/8-h night cycle. Twelve-day-old plants were inoculated with *Ph* or *Phb* and leaf samples were harvested at 14 dpi for DNA isolation, and 4 dpi (early time point) and 14 dpi (late time point) for RNA isolation, for a total of four samples. High-molecular-weight genomic DNA was isolated from *Ph*- or *Phb*-infected SusPtrit plants as previously described^33^. DNA integrity was confirmed using the FemtoPulse system (Agilent), and DNA concentration was estimated with the Qubit dsDNA HS Assay (Thermo Fisher Scientific). DNA purity was assessed by measuring 260/280 and 260/230 absorbance ratios with a Nanodrop spectrophotometer. Genomic DNA was sheared with the Megaruptor 3 (Diagenode, Denville, USA) to generate fragments of 15–20 kb. SMRTbell libraries were prepared using the SMRTbell prep kit 3.0 (102-182-700) and size-selected with the PippinHT system (Sage Science, HTP0001). Total RNA was extracted using the Maxwell RSC Plant RNA Kit (AS1500) on a Maxwell RSC48 instrument (Promega) according to the manufacturer’s protocol. RNA integrity was evaluated with an Agilent Bioanalyzer 2100, and the RNA Integrity Number (RIN) was ≥7.0 for all samples. RNA quantification was performed with the Qubit RNA HS Assay (Thermo Fisher Scientific). For Iso-Seq library preparation, ∼300 ng of high-quality total RNA was processed using the Kinnex full-length RNA kit (103-072-000; PacBio) following the recommended protocol. Final libraries QC were assessed with Qubit dsDNA High Sensitivity (ThermoFisher Scientific Q33230) and FEMTO Pulse (Agilent Technologies, Inc. P-0003-0817). For sequencing, SMRTbell libraries were prepared with the Revio polymerase kit (102-817-600) under conditions specified in SMRTlink. Sequencing was carried out on the PacBio Revio platform using the Revio sequencing plate (102-587-400) and Revio SMRT Cell Tray (102-202-200). DNA HiFi and Iso-Seq runs were performed in adaptive loading mode with movie times of 30 h and 24 h, respectively.

The raw genomics Revio reads from *Ph-* and *Phb*-infected barley were independently assembled using hifiasm (v0.19.8)^34^ with default parameters. Assembly qualities were assessed using the Quality Assessment Tool for Genome Assemblies (QUAST, v5.2) ^35^. The assembly metrics can be found in Supplementary Table 5. The resulting assemblies were filtered from the barley-specific contigs using IBSpy (v0.4.6)^36^. First, 31-mers sets were counted using KMC (v3.1.2)^37^ for GenBank assemblies GCA_038069595.1, GCA_949783145.1 and GCA_903970715.1 of barley and for GenBank assembly GCA_007896445.1 of *P. hordei 560*. The variation scores from each run using 50 kb window sizes against the two genome assemblies were independently stacked. The variation scores were manually inspected, *Puccinia*-specific contigs were extracted only if they showed high variation scores for the barley *k*-mer sets and low variation scores against the *Puccinia k*-mer sets. No hybrid *Ph*- or *Phb*-barley contigs were detected. The two filtered assemblies were inspected using BUSCO (v5.8.3)^38^ against the dataset pucciniomycetes_odb12. The Iso-Seq data were processed following the Iso-Seq3 pipeline (v4.0.0; https://github.com/PacificBiosciences/IsoSeq) separately for the four datasets from *Ph* and *Phb*-infected barley. In brief, after trimming the reads and removing the adapters with the lima tool (v2.12.0); the full-length non-chimeric reads (flnc) were refined using the Iso-Seq refine function. The next step was the generation of the consensus isoforms by using the Iso-Seq cluster function. RepeatMasker (v4.1.8)^39^ was used with the complete TREP database (v2019)^40^ followed by seqkit (v2.9.0) sort, to sort by size of the transcripts inside the isoform files. The resulting assemblies were filtered by mapping the isoforms over the respective filtered genomes using Minimap2 (v2.24)^41^ with the recommended setting for assembled isoform mapping (-ax splice:hq -uf). Successfully mapped isoforms were extracted using Samtools (v1.16.1)^42^.

### Genome and transcriptome sequencing of barley cultivar Vada

The experimental details and the relevant methods for barley cultivar Vada genome and transcriptome sequencing are described in Supplementary Note 1.

### Plasmid construction for expression in N. benthamiana and E. coli

For transient expression, genomic sequences of *Hv-LecRK, Hv-LecRK*^*SusPtrit*^ and *Hb*-*LecRK* including 2 kb of their upstream promoter regions, were synthesized in the entry vector pENTR/dTOPO (GenScript Biotech Corporation) and cloned into the modified version of the binary vector p6id35STE9 containing a Gateway cloning cassette and a FLAG-tag via Gateway LR reaction (Invitrogen). For domain swap constructs, the lectin domains of Hb-LecRK and Hv-LecRK were exchanged, and the resulting chimeric sequences were synthesized (GenScript Biotech Corporation). These constructs were expressed following the procedure described above. Rice *OsLecRK* sequences were retrieved from Rice Genome Annotation Project (http://rice.plantbiology.msu.edu/)^43^ and cloned using the same strategy.

For transient expression of candidate ligands, coding sequence (CDS) of 37 putative ligands were synthesized in the pENTR/dTOPO vector (GenScript Biotech Corporation) and cloned into the pGWB17 vector via Gateway LR reaction (Invitrogen). Thirty constructs with amino acid swap between *GH5*^*Ph1*^ and *GH5*^*Phb_h1*^ were synthesized by GenScript Biotech Corporation and cloned using the same strategy. For transient expression and co-immunoprecipitation (Co-IP) assays in *N. benthamiana*, the CDS of *GH5*^*Ph*^, *GH5*^*Phb_h1*^, *GH5*^*Phb_h2*^, *GH5*^*Pt*^, *GH5*^*Pgt*^, *GH5*^*Pst*^ and mutated *GH5*^*Ph*^ version (*mGH5*^*Ph*^) without stop codon were synthesized in the pENTR/dTOPO vector (GenScript Biotech Corporation) and cloned into the pGWB20 vector via Gateway LR reaction (Invitrogen). For quantitative Co-IP based on HiBit detection, all *GH5s* coding sequences were fused to a HiBiT tag and cloned into the pGWB20 vector.

For Co-IP assays, the CDS of *Hv-LecRK Hb*-*LecRK* and *OsLecRK-6* without stop codon were synthesized in the pENTR/dTOPO vector (GenScript Biotech Corporation) and cloned into the pGWB5 vector via Gateway LR reaction (Invitrogen). For protein expressions in *E. coli*, the codon optimized CDS without signal peptide of *GH5*^*Ph*^, *GH5*^*Phb*_*h1*^, *GH5*^*Phb*_*h2*^ and *m3GH5*^*Ph*^ were synthesized and cloned in the pJEX411C vector (Twist Biosciences). All constructs were sequence-verified prior to use. A list of plasmids and primers used in this study is listed in Supplementary Table 6 and 7, respectively.

### Agrobacterium-mediated transient expression in N. benthamiana

Three-to four-week-old *N. benthamiana* plants were thoroughly watered before infiltration. *A. tumefaciens* strains carrying the constructs of interest and the p19 RNAi suppressor were grown overnight at 28 °C in Luria–Bertani (LB) broth supplemented with appropriate antibiotics. Cells were pelleted by centrifugation at 3,000 rpm for 10 min and resuspended in infiltration buffer (10 mM MES pH 5.6, 10 mM MgCl_2_, 150 µM acetosyringone). The optical density at 600 nm (OD_600_) was adjusted as required, and suspensions were incubated at room temperature for 3 h prior to infiltration.

*A. tumefaciens* cultures carrying binary vectors were mixed with the p19 strain at a 1:1 ratio such that the final OD_600_ values were equal to 1.0 for constructs expressed under native promoters, and 0.5 for constructs expressed under the constitutive 35S promoter. For co-infiltration experiments, *A. tumefaciens* cultures were mixed with the p19 strain and adjusted to reach a final OD_600_ equal to 0.5. For constructs expressing *LecRKs* under native promoters, the OD_600_ was adjusted to 1.0, and for constructs expressing putative ligands under the constitutive 35S promoter, the OD_600_ was adjusted to 0.5. The mixtures were syringe-infiltrated into the abaxial side of *N. benthamiana* leaves. For cell death assays, leaves were collected 3 days post-infiltration (dpi), whereas for western blotting and Co-IP analyses, leaves were harvested 36 h and 30 h post-infiltration respectively.

### Apoplastic wash fluid extraction from barley leaves

Apoplastic wash fluid (AWF) was isolated from barley line SusPtrit, either uninfected (mock control) or inoculated with *Ph, Phb, Pt, Pst*, or *Pgt*, using the vacuum infiltration–centrifugation (VIC) method^44,45^. Approximately 25 g of fresh leaf tissue was harvested, leaves were cut into fragments and submerged in isolation buffer (0.05 M sodium phosphate, 50 mM Tris-HCl, pH 7.5, Milli-Q water). Vacuum was applied in 10 min intervals until tissues were fully saturated. Leaves were blotted dry, wrapped in parafilm, and placed in 50 ml Falcon tubes. Samples were centrifuged at 1,000 g for 15 min at 4 °C. The resulting AWF was collected and filtered through a 0.45 µm filter to remove the cellular debris and then snap-frozen in liquid nitrogen and stored at −80 °C until further use.

### AWF treatment and infiltration

Three days after the *A. tumefaciens* infiltration, 100 μl of apoplastic fluid was infiltrated into leaves at specific positions to evaluate hypersensitive response (HR). HR is a hallmark of plant disease resistance, and it has been shown that Hv-LecRK and Hb-LecRK trigger HR^6^. For heat treatment of AWF, samples were incubated at 94 °C for 15 min, immediately cooled on ice for 10 min, and centrifuged at 12,000 g for 10 min at 4 °C to remove precipitated material. The clarified supernatant was collected and used for infiltrations. For protease treatment of AWF, samples were treated with proteinase K at a final concentration of 100 µg/ml in buffer (50 mM Tris-HCl, pH 7.5) for 30 min at 37 °C. Reactions were terminated by addition of PMSF to 1 mM final concentration and incubation on ice for 10 min. Samples were subsequently cooled and used immediately for downstream infiltration and cell death assays. For lipase treatment, aliquots of AWF were incubated with lipase from *Aspergillus niger* (200 U/g) at final activities of 0.5-1.0 U/ml (equivalent to 2.5-5 mg/ml) in the same buffer for 30 min at 37 °C. Reactions were terminated by ultrafiltration through 10 kDa MWCO spin columns to remove enzyme.

### HR imaging in Vilber FUSION FX

The leaf imaging was performed two days after apoplastic wash fluid infiltration and three days after co-infiltration experiments. Leaf discs (0.5 mm in diameter) corresponding to each infiltration site were collected using a cork-borer, and imaged at a wavelength of 480 nm using a Vilber FUSION FX Imaging system (www.vilber.com) (Vilber Lourmat, Eberhardzell, Germany) with the following settings: fluorescence sample; excitation: blue epi-illumination; emission: filter F-535 Y2; aperture: 0.84-open; sensitivity: full resolution. Later, the images were analyzed for HR quantification using Image J software (https://imagej.nih.gov/ij/)^46^. The mean values for HR intensity were normalized to the infiltrated leaf area.

### Sample preparation and LC-MS/MS of AWF protein extracts

AWF proteins were precipitated using tri-chloro acetic acid (TCA)/acetone^47^. The precipitated proteins were resuspended in 100 µl of 8 M urea in 100 mM triethylammonium bicarbonate (TEAB, pH 8.5; Sigma-Aldrich, T7408).

Proteins were reduced with 10 mM tris (2-carboxyethyl) phosphine (TCEP, Sigma-Aldrich, C-4706) at 37 °C for 1 h and alkylated with 20 mM methyl methanethiosulfonate (MMTS, Sigma-Aldrich) at room temperature for 15 min. Samples were diluted to 1 M urea with 100 mM TEAB and digested overnight at 37 °C with sequencing-grade modified porcine trypsin (Promega, 1:50 enzyme:substrate, w/w). Digestion was stopped by adding formic acid (FA) to 1%, and peptides were desalted using C18 ZipTips (Millipore, ZTC18S096), dried in a SpeedVac, and resuspended for LC-MS/MS analysis. All solutions were prepared with ultrapure water (Milli-Q, Millipore). LC-MS/MS was performed on an Orbitrap Fusion Lumos mass spectrometer coupled to a Dionex UltiMate 3000 RSLC system (Thermo Scientific). Peptides were trapped on a PepMap NEO C18 column (300 µm × 5 mm, 5 µm) and separated on a PepMap RSLC C18 analytical column (75 µm × 50 cm, 2 µm, 100 Å) at 300 nl/min. Mobile phases were water with 0.1% FA (solvent A) and 95% acetonitrile (ACN) with 0.1% FA (solvent B). Peptides were eluted using a 59 min gradient: 2–6% B for 2 min, 6–31.6% B over 52 min, 31.6–95% B over 5 min, followed by 10 min at 95% B and re-equilibration at 2% B. The EASY-Spray emitter was operated at 1900 V in positive ion mode with lock mass m/z 445.120025. Survey scans (m/z 350–1500) were acquired at 60,000 resolutions, AGC target 4.0 × 10^5^, and maximum injection time 50 ms. The most intense ions (charge 2+ to 5+/6+) were fragmented by higher-energy collisional dissociation (HCD) at 30% collision energy, isolated with a 1.6 Th window, AGC target 5.0 × 10^4^, and dynamic max injection time. Fragment spectra were recorded at 30,000 resolutions.

Raw data were processed using MASCOT v2.7 (Matrix Science). Protein identification was performed against the barley cv. Vada database (generated in this study) (Supplementary Note 1) and the *P. hordei Ph560* database^12^. Search parameters included trypsin with up to two missed cleavages, carbamido-methylation of cysteine as a fixed modification, 10 ppm precursor mass tolerance, and 0.02 Da fragment ion tolerance. Peptide-spectrum matches (PSMs), and protein identifications were filtered at a 1% false discovery rate (FDR). The barley cv. Vada database was used to remove plant proteins, while *Ph560* was used to identify fungal proteins. A total of 232 fungal proteins were identified from the AWF, consistently detected across five independent biological replicates, each protein with at least three unique peptides.

### AlphaFold-based screening of candidate ligands

The lectin domains of Hv-LecRK (residues 25–290) and Hb-LecRK (residues 32–290) were used for interaction analyses. Protein–ligand interactions were predicted against a set of 232 candidate ligands using a customized implementation of AlphaFold-Multimer^48,49^ on the Ibex high-performance computing cluster at KAUST. This implementation decouples feature generation from inference, such that multiple-sequence alignments and structural features for each protein chain are generated once on CPU nodes and subsequently re-used during GPU-based inference. For each predicted complex, the ipTM+pTM composite scores (default AlphaFold-Multimer quality score) and the ipTM scores were calculated. The ipTM score, which specifically evaluates confidence in the predicted protein–protein interface, is more robust to the presence of disordered regions in non-interacting protein segments. The quality scores for all predicted interactions of Hv-LecRK and Hb-LecRK lectin domains with the 232 candidate ligands are provided in Supplementary Table 1 and 2.

### Protein expression and purification from E.coli and N. benthamiana AWF

Recombinant constructs encoding GH5^Ph^, GH5^Phb_h1^, and GH5^Phb_h2^, each with an N-terminal His-tag were expressed in *E. coli* Rosetta (DE3) cells. Cultures were grown in LB medium at 37° C until reaching an optical density at OD_600_ equal to 0.6. Protein expression was induced with 0.2 mM isopropyl β-D-1-thiogalactopyranoside (IPTG), followed by overnight incubation at 16 °C to promote soluble expression. Cells were harvested and lysed by sonication in buffer A (50 mM Tris-HCl, pH 7.5; 300 mM NaCl; 10 mM imidazole; 1 mM dithiothreitol [DTT]; 1% Triton X-100). Lysates were clarified by centrifugation at 75,000 × g for 30 min at 4 °C. The supernatant was filtered and loaded onto a Ni-NTA affinity column (Cytiva) pre-equilibrated with buffer A, followed by washing with 15 column volumes of the same buffer. Bound proteins were eluted using buffer B (50 mM Tris-HCl, pH 7.5; 300 mM NaCl; 500 mM imidazole; 1 mM DTT; 1% Triton X-100). Eluted proteins were further purified by size-exclusion chromatography using a Superdex 200 16/600 column (Cytiva) equilibrated with buffer containing 20 mM HEPES (pH 7.5), 150 mM NaCl, and 3 mM DTT. Target proteins eluted as distinct peaks corresponding to their expected molecular weights, confirming successful purification and homogeneity.

GH5^Ph^, GH5^Phb_h1^, and GH5^Phb_h2^ proteins were expressed and purified from the *N. benthamiana* apoplast as previously described^50^ with slight modifications. An N-terminal PR-1a signal peptide was fused to GH5 proteins to facilitate efficient secretion into the *N. benthamiana* apoplast, and C-terminal HiBiT and 6XHis tags were incorporated for quantitative HiBiT detection and affinity purification, respectively. The resulting constructs were cloned into the pSuper1300 expression vector and transiently expressed in *N. benthamiana* leaves via *A. tumefaciens*-mediated infiltration. Agrobacterium-infiltrated plants were incubated for five days. For AWF protein extraction, 60-70 leaves were harvested and subjected to vacuum infiltration with infiltration buffer (20 mM Bis-Tris/HCl (pH 6.0), 0.01% (v/v) Tween-20, and 1× cOmplete™ Protease Inhibitor Cocktail), followed by centrifugation-based apoplastic fluid collection as described previously. For purification, clarified AWF extracts were subjected to Ni^2+^-NTA affinity chromatography using binding buffer (20 mM Bis-Tris/HCl, pH 6.0, 300 mM NaCl). The resin was washed extensively with wash buffer containing 10 mM imidazole to remove non-specifically bound proteins, and His-tagged proteins were eluted using buffer containing 300 mM imidazole. Eluted fractions were analyzed by SDS–PAGE followed by Coomassie staining. Target proteins correspond to their expected molecular weights, confirming successful purification and homogeneity. Protein concentration was determined by absorbance at 280 nm. Purified proteins were concentrated and buffer-exchanged using a 10 kDa molecular weight cut-off ultrafiltration device as required.

### Enzymatic assay

The enzymatic activity of GH5 proteins was assayed using pustulan (Biosynth, YP15423), as a substrate^51,52^.The reaction mixture (100 µl) contained 1 µg ml^−1^ purified GH5 protein and 0.9% (w/v) pustulan in either 50 mM sodium acetate buffer (pH 5.5) or 50 mM ammonium acetate buffer (pH 5.5) were incubated at 30°C with shaking at 450 rpm for 3 h for glucose quantification and 1h for gentiobiose detection. Pustulanase (Prokazyme, Cel136) was used as a positive control at a final activity of 1.0 U ml^−1^ under the same reaction conditions. Reactions were terminated by boiling for 5 min, followed by centrifugation at 16,000g for 2 min to remove denatured proteins. The resulting supernatants were collected for downstream analyses. Glucose release in the supernatant was quantified using a glucose oxidase/peroxidase-based Glucose Assay Kit (Abcam, ab65333) according to the manufacturer’s instructions.

For gentiobiose detection, supernatants from reactions prepared in ammonium acetate buffer were analyzed by LC–MS using a UHPLC-Q-Orbitrap-MS system (Q-Exactive Plus) operated in positive ionization mode with full-scan acquisition. Instrument parameters were set as follows: spray voltage, 3.5 kV; capillary temperature, 250 °C; auxiliary gas temperature, 310 °C; sheath gas, 30 Arb; and auxiliary gas, 10 Arb. UHPLC separation was performed on an UltiMate 3000 UHPLC system (Thermo Fisher Scientific) equipped with a BEH HILIC column (Waters; 2.1 × 100 mm, 1.7 µm) maintained at 30 °C. The mobile phases consisted of water containing 0.1% (v/v) formic acid (solvent A) and acetonitrile containing 0.1% (v/v) formic acid (solvent B). Separation was carried out at a flow rate of 0.2 ml min^−1^ using the following gradient: 1% B for 5 min, increased to 100% B over 10 min, held at 100% B for 5 min, and re-equilibrated at 1% B for 4 min. Gentiobiose was quantified using QuanBrowser software based on the potassium adduct ion [M+K]^+^ at *m/z* 381.0793.

### Protein extraction and immunoblotting

*N. benthamiana* leaves were harvested, and ground into fine powder using liquid nitrogen, and the total proteins were solubilized with the extraction buffer (50 mM Tris-HCl, pH 7.5, 150 mM NaCl,1 mM EDTA, 0.5% Triton X-100, 10% glycerol, 2 mM DTT, 1 mM PMSF and protease inhibitor cocktail, Roche). The samples were mixed by rotating at 4 °C for 15 minutes, then centrifuged at 4 °C for 25 min at 16,000 x g and the supernatant was collected. Approximately 20 μg from each protein sample were separated by SDS-PAGE and transferred to 0.22 μM polyvinylidene fluoride (PVDF) membrane (Immobilon, Merck Millipore). Protein loading was visualized by staining membranes with Ponceau S (Thermo Scientific). Membranes were blocked with EveryBlot Blocking Buffer (Biorad) and then incubated in the primary and secondary antibodies. Blots were probed with HRP-conjugated rabbit anti-DDDDK tag (binds to FLAG tag sequence) antibody (1:5000, abcam, ab1162) or rabbit anti-MYC (1:1000, Cell Signaling Technology, 2278) and followed by goat anti-rabbit antibodies conjugated with horseradish peroxidase (1:15000, Invitrogen), to detect FLAG and MYC-tagged proteins, respectively. Protein signal detection was performed by the ECL detection kit (Cytiva; RPN2235) and recorded using Vilber FUSION FX Imaging system.

### Co-immunoprecipitation

Co-IP experiments were performed as previously described with minor modifications^53^. *N. benthamiana* leaves were harvested and ground to a fine powder using liquid nitrogen. Proteins were extracted in 1 ml of extraction buffer (50 mM Tris-HCl, pH 7.5, 150 mM NaCl, 1 mM EDTA, 10% glycerol, 1 mM PMSF, 5 mM DTT, 0.5% NP-40, and protease inhibitor cocktail, Roche). The samples were mixed by rotating at 4 °C for 15 minutes, then centrifuged at 4 °C for 25 min at 16,000 x g and the supernatant was collected. A total of 800 μl of the supernatant was incubated with 10 μl pre-equilibrated GFP-Trap Magnetic Agarose beads (Chromotek) for 90 min at 4 °C with gentle rotation. Beads were washed once with extraction buffer and four times with wash buffer (50 mM Tris-HCl, pH 7.5, 150 mM NaCl, 1 mM EDTA). Co-immunoprecipitated proteins were then detected via western blot using anti-GFP (Cell Signaling Technology, 2956) or anti-MYC (Cell Signaling Technology, 2278) antibodies.

NanoLuc-based HiBiT–LgBiT complementation system was used to perform HiBiT–based quantitative co-IP^54,55^ using the same protein extraction and GFP-Trap immunoprecipitation procedure described above. Briefly, total proteins were extracted from *N. benthamiana* leaves and incubated with GFP-Trap Magnetic Agarose beads (Chromotek). Following washing, bead-bound proteins were eluted by boiling in elution buffer containing 100 mM Tris-HCl (pH 8.0) and 100 mM NaCl. For HiBiT-based quantification, 25 µl of the eluted samples were transferred to a 96-well plate pre-blocked with blocking buffer (BSA) for 20 min at room temperature. Subsequently, 75 µl of freshly prepared HiBiT detection reagent (Nano-Glo® HiBiT Blotting System kit, Promega™) was added to each well, and incubated for 10 min at RT. The luminescence was measured using a microplate reader (GloMax® Navigator Microplate Luminometer) with an integration time of 1 s.

### Binding assay

Quantitative binding assays based on the NanoLuc-based HiBiT–LgBiT complementation system were adapted from a previously described protocol^18^ with minor modifications, and all buffer compositions were prepared as described therein. GFP-tagged Hv-LecRK or Hb-LecRK proteins were transiently expressed in *N. benthamiana* leaves via *A. tumefaciens*-mediated infiltration. Leaf tissues were harvested 36 hours post infiltration, frozen in liquid nitrogen and ground to a fine powder. Total proteins were extracted in ice-cold extraction buffer and solubilized by incubation at 4 °C with gentle rotation. Following centrifugation at 16,000 × g for 25 min at 4 °C, the resulting supernatant was used for receptor immobilization.

For receptor immobilization, GFP-Trap Magnetic Agarose beads (Chromotek) were equilibrated in solubilization buffer and incubated with GFP-tagged LecRK-containing extracts for 1 h at 4 °C with gentle rotation. Beads were subsequently washed twice with solubilization buffer, twice with IP wash buffer and twice with binding buffer. Binding reactions were initiated by incubating immobilized GFP-tagged Hv-LecRK or Hb-LecRK or GFP only controls with a dilution series (0.0001–100 µM) of HiBiT-tagged GH5 proteins purified from *N. benthamiana* AWF, in binding buffer for 45 min at 4 °C with gentle rotation. Following binding, beads were washed four times with binding wash buffer to remove unbound proteins.

For quantification, bead-bound proteins were resuspended in elution buffer and boiled at 95 °C for 10 min to release bound HiBiT-tagged GH5 proteins. Eluted samples (10 µl) were transferred to BSA-blocked 96-well plates, followed by addition of 90 µl freshly prepared HiBiT detection reagent (Nano-Glo® HiBiT Blotting System kit, Promega™) to each well, and incubated for 10 min at RT. Luminescence was measured using a GloMax® Navigator microplate luminometer with an integration time of 1 s. Relative luminescence units (RLU) were used as a quantitative measure of GH5–LecRK binding. Background luminescence was determined using control reactions containing GFP-Trap beads incubated with HiBiT-tagged GH5 proteins in the absence of immobilized receptor proteins, and these values were subtracted from all measurements before analysis. For binding curve analysis, background-subtracted signals were normalized to maximal binding obtained at saturating ligand concentrations. Dissociation constants (K_d_) were determined by nonlinear regression using a one-site specific binding model in GraphPad Prism (GraphPad Software).

### M. oryzae transformation

The CDS of *GH5*^*Ph*^, *GH5*^*Phb_h1*^ and *GH5*^*Phb_h2*^ and *m3GH5*^*Ph*^ variants were cloned with *RP27* or *PWL2* promoter with *TrpC* terminator into the pCB1532H-RFP vector (pCB1532H-RFP was a gift from Sophien Kamoun, Addgene plasmid, 101855) (GenScript Biotech Corporation)^56^. Promoter sequences for *RP27* and *PWL2* were obtained from *M. oryzae*, and the *TrpC* terminator sequence was derived from *Aspergillus nidulans*. The resulting constructs were introduced individually into the wild-type *M. oryzae* strain FR13 by protoplast-mediated transformation as previously described^57^. The transformants were selected on medium supplemented with hygromycin B (200 μg/ml). Genomic DNA was extracted from *M. oryzae* transformants following established protocol^32^. Transformants were verified by PCR using gene-specific primers. The details of all constructs used for transformation are provided in Supplementary Table 6 and primer sequences are listed in Supplementary Table 7.

### Wheat transformation

A construct containing the genomic sequence of *Hv-LecRK* or *Hb-LecRK* with native promoter was transformed into wheat cv. Fielder as a service by the Wisconsin Crop Innovation Center (WCIC), University of Wisconsin–Madison. Two independent homozygous lines (E1 and E2) were selected for phenotypic evaluation in growth chamber and field conditions. For wheat transformation with *Os-LecRK*, the genomic sequence of *OsLecRK-6* and 2 kb of its upstream promoter region was synthesized and cloned into an entry pENTR/dTOPO vector (GenScript Biotech Corporation) (Supplementary Table 6). The entry construct was then introduced into the modified version of the binary vector p6id35STE9 containing a Gateway cloning cassette and a FLAG-tag via Gateway LR reaction (Invitrogen). The resulting construct was used for *A. tumefaciens*-mediated transformation of wheat cultivar Fielder. Stable wheat transgenic plants were generated as previously described^58^. Two independent *OsLecRK-6* T0 lines were obtained (E1 and E2). The positive transformants were confirmed by PCR using gene-specific primers (Supplementary Table 7). Homozygous T2 plants for *Os-LecRK-6* were selected for phenotypic evaluation.

### Field experiments

Leaf rust field experiments were conducted during the 2023–2024 and 2024–2025 growing seasons at KAUST Agricultural Research Field Site. Wheat transgenic lines *Hv-LecRK-E1, Hv-LecRK-E2, Hb-LecRK-E1, Hb-LecRK-E2*, and their corresponding null segregants were evaluated under field conditions. Lines were planted in 1.0 m rows containing 15 plants per row, with four replicates arranged in a randomized design. The experimental field was surrounded by border spreader rows and within the field, transgenic and null segregant plots were alternated and interspersed with spreader rows to ensure uniform disease pressure. Spreader rows consisted of a 1:1:1 mixture of highly susceptible wheat cultivars ‘Thatcher’, ‘Fielder’, and ‘Arina’ to maintain high and uniform *Pt* inoculum pressure across the field. Sowing was performed on December 21^st^ for both field seasons (2023–2024 and 2024–2025). Spreader rows were inoculated with a mixture of seven *Pt* isolates as previously described^59^; details of *Pt* isolates are provided in Supplementary Table 4. Artificial inoculation was initiated at 30–31 days after sowing (January 21^st^, 2024 and January 20^th^, 2025) by introducing urediniospore-infected plants into spreader rows, followed by two weekly spray inoculations of a urediniospore suspension onto spreader rows to ensure uniform disease establishment. Repeated leaf rust observations were made throughout the growing season. Leaf rust severity was visually estimated as the percentage of leaf area covered by rust pustules, using the modified Cobb scale^60^. Disease progression on flag leaves was assessed at three time points for each field season. Final leaf rust severity on the flag leaves was recorded when susceptible null segregant plants reached 60% or more, ensuring saturation of infection pressure.

### Phylogenetic analysis of GH5 members

All the protein sequences corresponding to the GH5 family for seven different rust pathogens were extracted from the Carbohydrate Active Enzymes database (CAZy) website (http://www.cazy.org/)^61^ (Supplementary Table 8). Multiple sequence alignment of these proteins was performed using Clustalw2.1^62^. In parallel, InterProScan (v5.64-96.0)^63^ with default parameters for the following databases: FunFam, SFLD, PANTHER, Gene3D, PRINTS, Coils, SUPERFAMILY, SMART, CDD, PIRSR, ProSitePatterns, AntiFam, Pfam, MobiDBLite, PIRSF, NCBIfam, was run locally to confirm the enzyme classification for each of the retrieved proteins. Both the MSA and the InterProScan output were fed into a Python script to produce the final phylogenetic tree. OpenAI’s ChatGPT (https://chat.openai.com) was used to assist the generation of the script for the phylogenetic tree.

### Statistical analysis

All statistical analyses were performed using two-sample Student’s *t*-Test or Wilcoxon Signed-Rank Test for paired samples. Data were first tested for normality (Shapiro-Wilk test) and the corresponding paired tests were applied. Statistical significance was reported using two-tailed *p*-values, which are indicated in each figure, above the comparisons. Data analysis was generated using the Real Statistics Resource Pack software (Release 8.9.1), Copyright (2013-2023) Charles Zaiontz (www.real-statistics.com), the Analysis ToolPak from Microsoft Excel (2025. Microsoft Corporation); and BioRender Graph (BioRender.com). The number of biological replicates for each experiment is indicated in the corresponding figure legends.

## Data availability

The main data supporting the findings of this study are available within the article, its Extended Data Figures and its Supplementary Information. For more details contact the corresponding authors.

