## Extended Data Figures for "Host specificity in cereal rust fungi is mediated by a conserved glycoside hydrolase family"

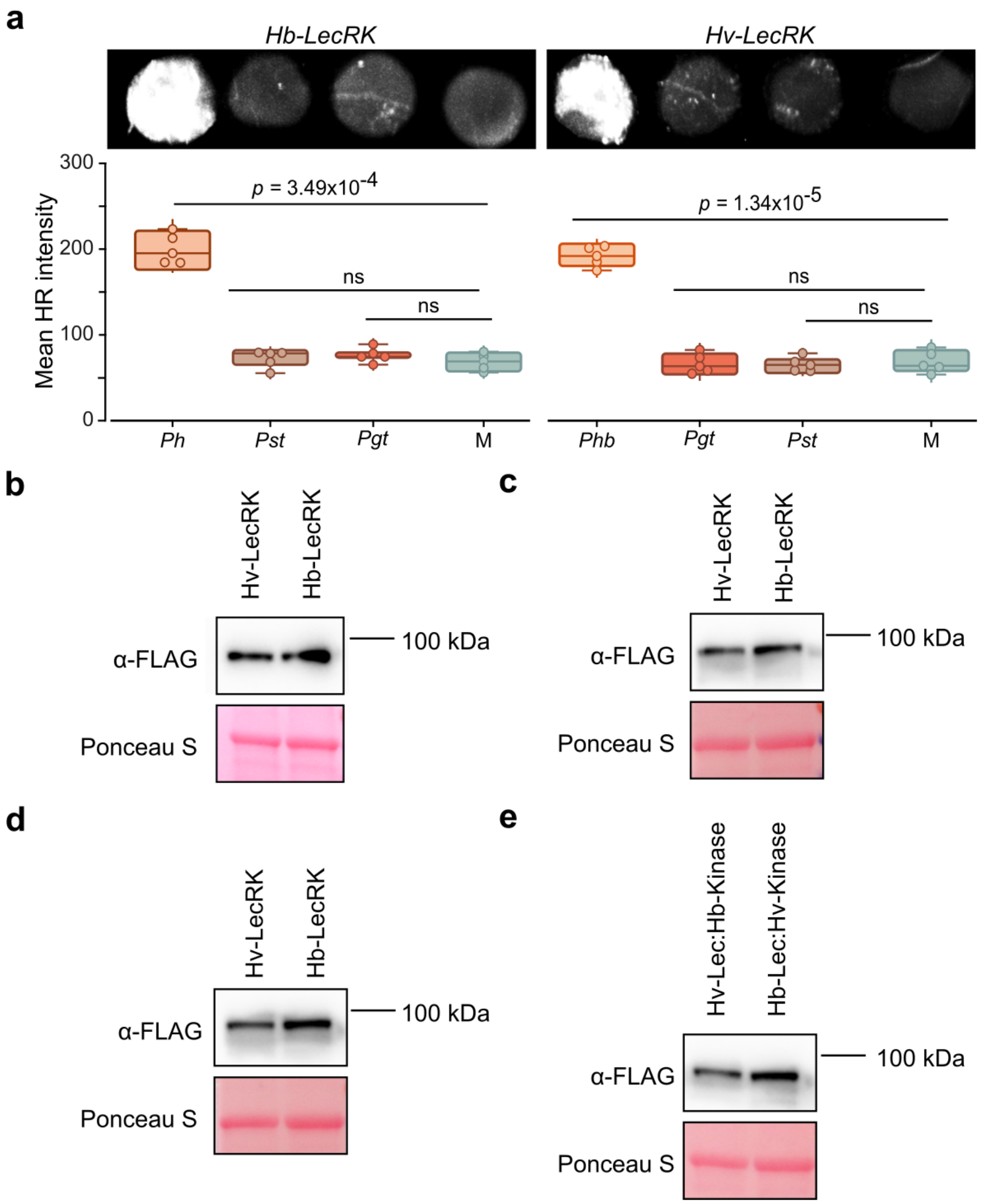


**Extended Data Fig. 1. Apoplastic wash fluid (AWF) from *Pgt* and *Pst*-infected barley leaves failed to induce HR. a**, Transient expression of *Hb-LecRK* and *Hv-LecRK* in *N. benthamiana* followed by infiltration with apoplastic wash fluid (AWF) derived from *Pgt*- and *Pst*-infected barley. HR was imaged (top) two days after AWF infiltration (Fusion-FX imager) and quantified (bottom) using ImageJ. Box plots represent the quantification of mean HR intensity (dark orange, *Ph*; light orange, *Phb*; dark brown, *Pst;* salmon, *Pgt*) of five biological replicates (*n* = 5). Circles indicate individual data points. The horizontal line denotes the median, box edges mark the first (Q1) and third (Q3) quartiles, and whiskers extend to the minimum and maximum values**.** Statistical significance was determined by paired two-sample Student's *t*-Test or by Wilcoxon Signed-Rank Test for paired samples; *p*-values are indicated above the comparisons. Cases with no significant difference are indicated with "ns". The experiments were repeated three times with consistent results. **b**, Immunoblots showing protein abundance of Hb-LecRK and Hv-LecRK. **c**, Immunoblots showing protein abundance of Hv-LecRK and Hb-LecRK for transient assay performed in Fig. 1b. **d**, Immunoblots showing protein abundance of Hv-LecRK and Hb-LecRK for transient assay performed in Fig. 1c. **e**, Immunoblots showing protein abundance of domain swap constructs encoding chimeric LecRK variants, Hv-Lec:Hb-Kinase and Hb-Lec:Hv-Kinase used in Fig. 1d. For **b-e**, *Hb-LecRK* and *Hv-LecRK*, and domain swap constructs were driven by their native promoters and were transiently expressed in *N. benthamiana* leaves. Total protein was extracted at 36 h post-infiltration and detected by anti-FLAG antibodies.

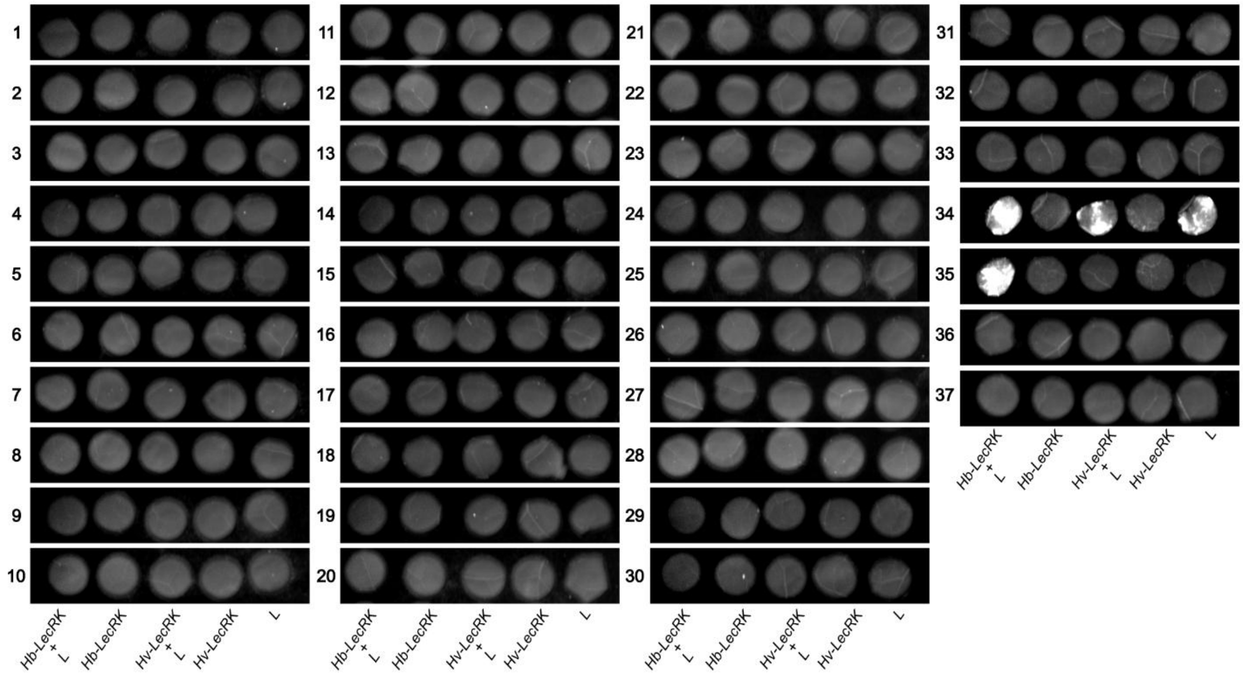


**Extended Data Fig. 2. Screening of 37 candidates from *Ph* using the *N. benthamiana* transient assay.** HR imaging in *N. benthamiana* co-expressing each candidate with *Hb-LecRK* or *Hv-LecRK*. HR was recorded three days post-infiltration (Fusion-FX imager). Candidate 34 induced an auto-active cell death response independent of Hb-LecRK. Candidate 35, $GH5^{Ph}$ from *Ph560*, triggered HR specifically in the presence of Hb-LecRK. 'L' stands for candidate ligand. Gene IDs for the different candidates are provided in Supplementary Table 3.

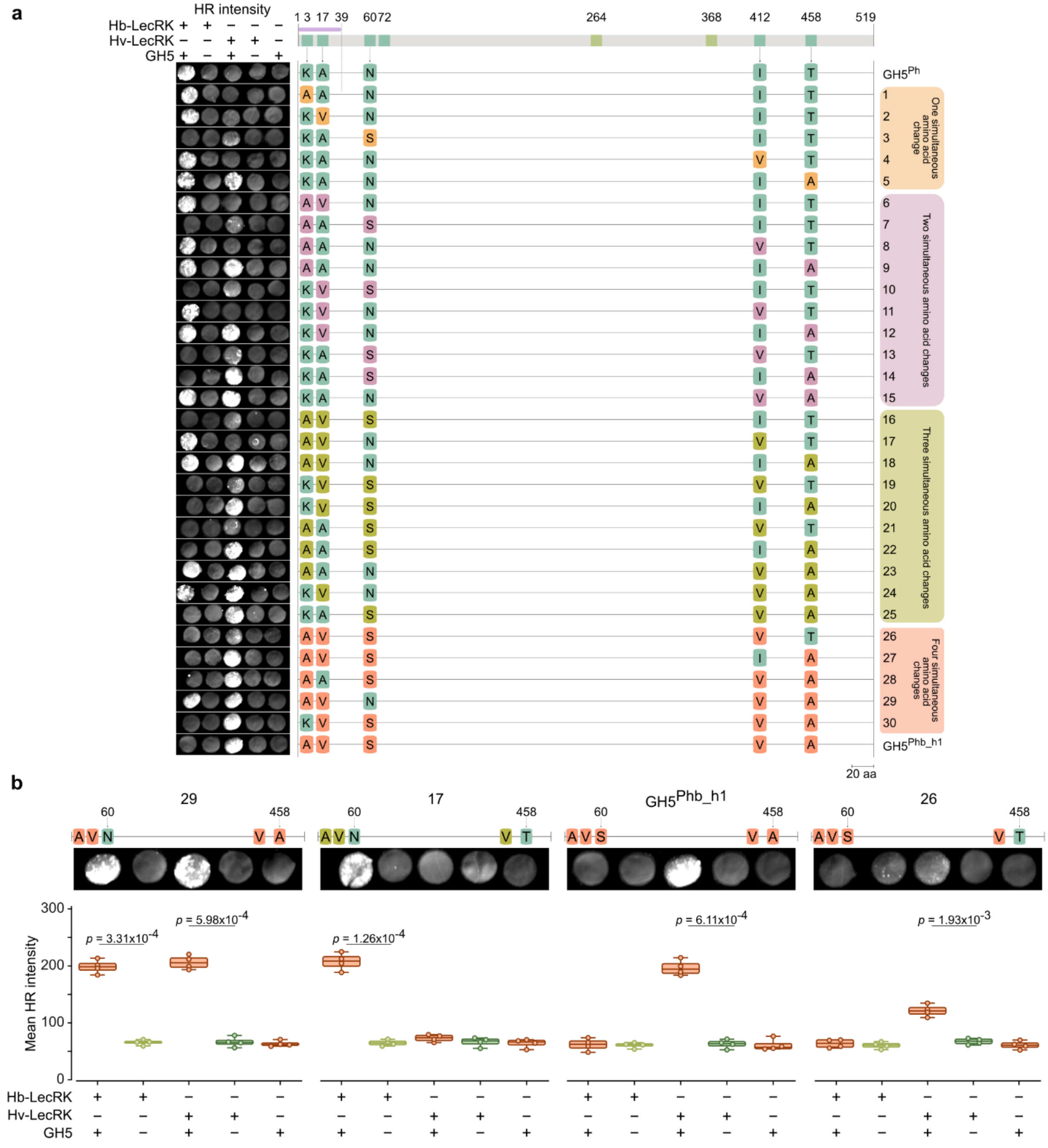


**Extended Data Fig. 3. Amino acid swaps between GH5$^{Ph}$ and GH5$^{Phb_h1}$ identify two residues that contribute to recognition specificity. a**, HR imaging in *N. benthamiana* leaves co-expressing *GH5$^{Ph}$* variants with *Hb-LecRK* or *Hv-LecRK*. A total of thirty GH5$^{Ph}$ variants were tested, each carrying substitutions at one (soft orange), two (mauve), three (olive) or four (salmon) polymorphic amino acid positions, in which residues were replaced with the corresponding amino acids from GH5$^{Phb_h1}$. **b**, HR imaging (top) and quantification (bottom) in *N. benthamiana* leaves co-expressing *GH5$^{Ph}$* variant 29, *GH5$^{Ph}$* variant 17, wild-type *GH5$^{Phb_h1}$* and *GH5$^{Ph}$* variant 26 with *Hb-LecRK* or *Hv-LecRK*. Corresponding *GH5* variants and *LecRKs* expressed alone are shown as controls. HR was imaged three days post-infiltration

(Fusion-FX imager). Box plots represent the quantification of mean HR intensity of ten biological replicates ($n$ = 10). Circles indicate individual data points. The horizontal line denotes the median, box edges mark the first (Q1) and third (Q3) quartiles, and whiskers extend to the minimum and maximum values**.** Statistical significance was determined by paired two-sample Student's $t$-Test; $p$-values are indicated above the comparisons.

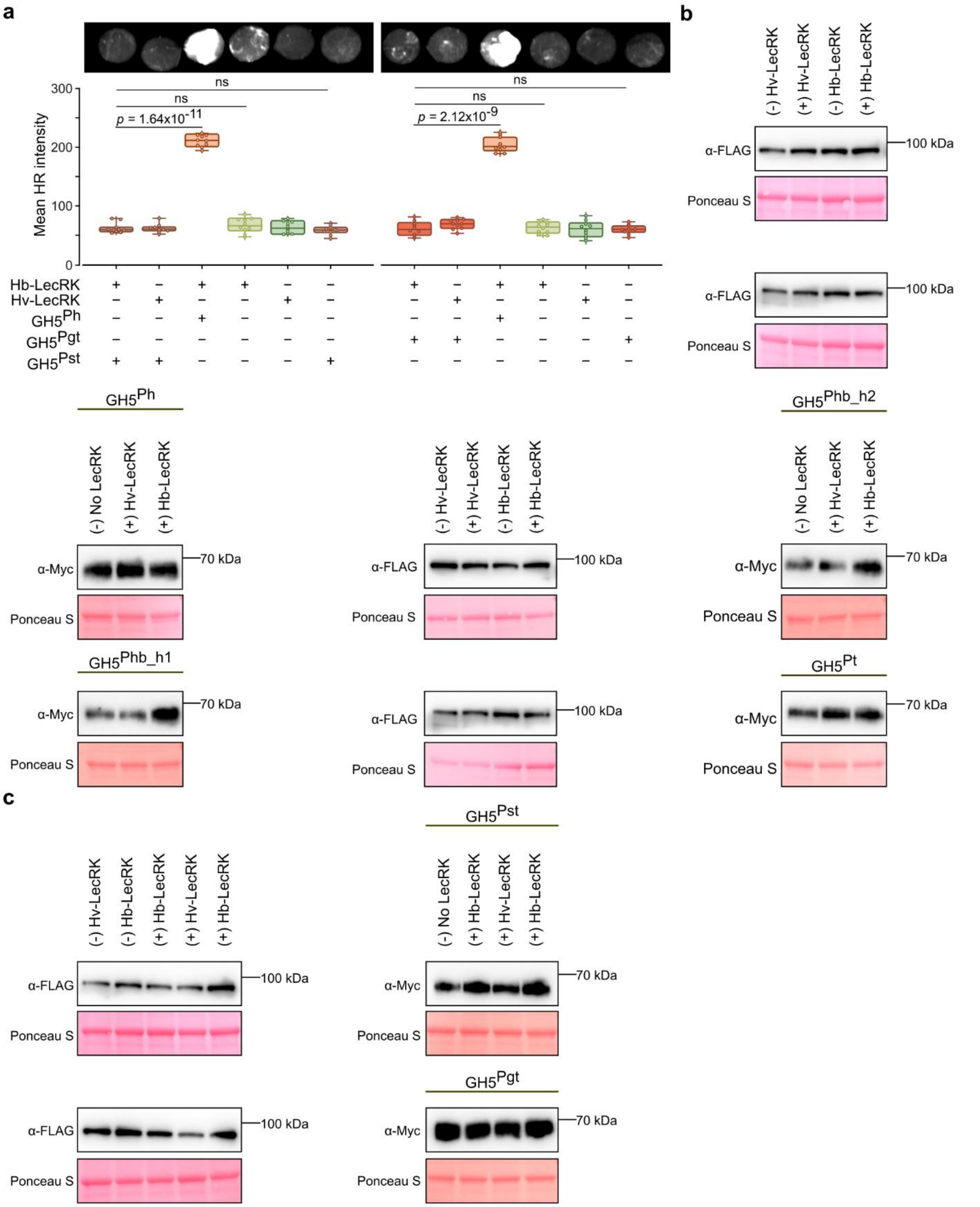


**Extended Data Fig. 4. *GH5^Pst* and *GH5^Pgt* failed to induce HR when co-expressed with *Hv-LecRK* and *Hb-LecRK*. a**, HR imaging (top) and quantification (bottom) in *N. benthamiana* co-expressing *GH5^Pst*,

*GH5$^{Pgt}$* and *GH5$^{Ph}$* with *Hb-LecRK* or *Hv-LecRK*. Corresponding *GH5s* and *LecRKs* expressed alone are shown as controls. HR was imaged three days post-infiltration (Fusion-FX imager). Box plots represent the quantification of mean HR intensity (light orange, GH5$^{Ph}$; dark brown, GH5$^{Pst}$; salmon, GH5$^{Pgt}$; expressed with *LecRKs* or alone; light green, *Hb-LecRK* alone; dark green, *Hv-LecRK* alone) of ten biological replicates ($n$ = 10). Circles indicate individual data points. The horizontal line denotes the median, box edges mark the first (Q1) and third (Q3) quartiles, and whiskers extend to the minimum and maximum values**.** Statistical significance was determined by paired two-sample Student's *t*-Test; *p*-values are indicated above the comparisons. Cases with no significant difference are indicated with "ns". **b**, Immunoblots showing protein abundance of Hv-LecRK, Hb-LecRK, GH5$^{Ph}$, GH5$^{Phb_h1}$, GH5$^{Phb_h2}$ and GH5$^{Pt}$ refers to Fig. 2b. **c**, Immunoblots showing protein abundance of Hv-LecRK, Hb-LecRK, GH5$^{Pst}$ and GH5$^{Pgt}$. The GH5 variants from different rust species were co-expressed with Hb-LecRK or Hv-LecRK in *N. benthamiana* leaves. (+) indicates co-expression of lectin receptor kinase with GH5, (-) indicates infiltration of lectin receptor kinase alone, 'No LecRK' indicates GH5 expressed alone. Total protein was extracted at 36 h post-infiltration and expression of Hv-LecRK and Hb-LecRK were detected by anti-FLAG antibodies. Expression of GH5$^{Ph}$, GH5$^{Phb_h2}$, GH5$^{Phb_h1}$, GH5$^{Pt}$, GH5$^{Pst}$ and GH5$^{Pgt}$ were detected by anti-Myc antibodies. The experiments were repeated three times with consistent results.

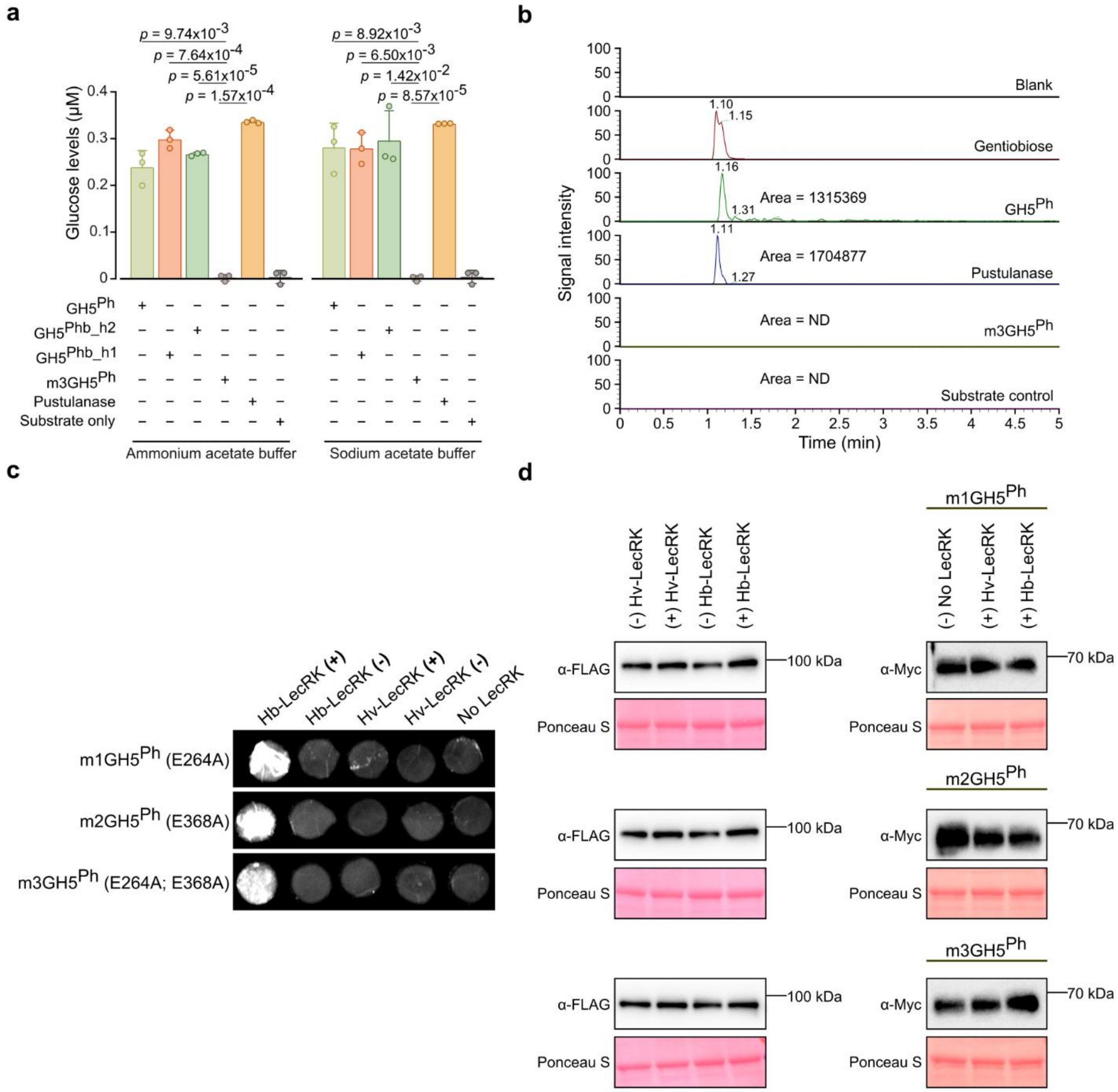


**Extended Data Fig. 5. Enzymatic activity of GH5 proteins and HR response of catalytic GH5 mutants.** **a**, Enzyme activity of GH5 proteins determined by quantifying glucose released from pustulan following incubation with $GH5^{Ph}$, $GH5^{Phb\_h1}$, $GH5^{Phb\_h2}$, catalytically inactive $GH5^{Ph}$ variants ($m3GH5^{Ph}$; E264A and E368A), or pustulanase (positive control) at 30°C in 50 mM sodium acetate or ammonium acetate buffer (pH 5.5). Substrate-only reactions served as negative controls. Data are presented as mean ± s.d. glucose concentrations from three independent replicates (n = 3). Circles indicate individual data points. The statistical significance of differences was determined by paired two-sample Student's *t*-tests (two-tailed); *P*-values are indicated above the comparisons. **b**, LC-MS analysis for detection of gentiobiose released from pustulan after incubation with $GH5^{Ph}$, $m3GH5^{Ph}$ or pustulanase (positive control) at 30°C in 50 mM ammonium acetate buffer (pH 5.5). Substrate-only reactions served as negative controls. Area under peak corresponds to relative abundance. **c**, HR imaging in *N. benthamiana* leaves co-expressing the catalytic mutant variant *GH5*$^{Ph}$ E264A (m1) with *Hb-LecRK* (top), *GH5*$^{Ph}$ variant E368A (m2) with *Hb-LecRK* (middle), and *GH5*$^{Ph}$ variant E264A and E368A (m3) with *Hb-LecRK* (bottom). HR was imaged three days post-infiltration using Fusion-FX imager. **d**, Immunoblots showing protein abundance of Hv-LecRK, Hb-LecRK and mutated versions of GH5$^{Ph}$ m1, m2 and m3. The GH5 variants from different rust species were co-expressed with Hb-LecRK or Hv-LecRK in *N. benthamiana* leaves. (+) indicates co-expression of lectin receptor kinase with GH5, (-) indicates infiltration of lectin receptor kinase alone, 'No LecRK' indicates GH5 expressed alone. Total protein was extracted at 36 h post-infiltration and expression of Hv-LecRK and Hb-LecRK were detected by anti-FLAG antibodies. Expression of mutated version of GH5$^{Ph}$ were detected by anti-Myc antibodies. All experiments were repeated three times with consistent results.

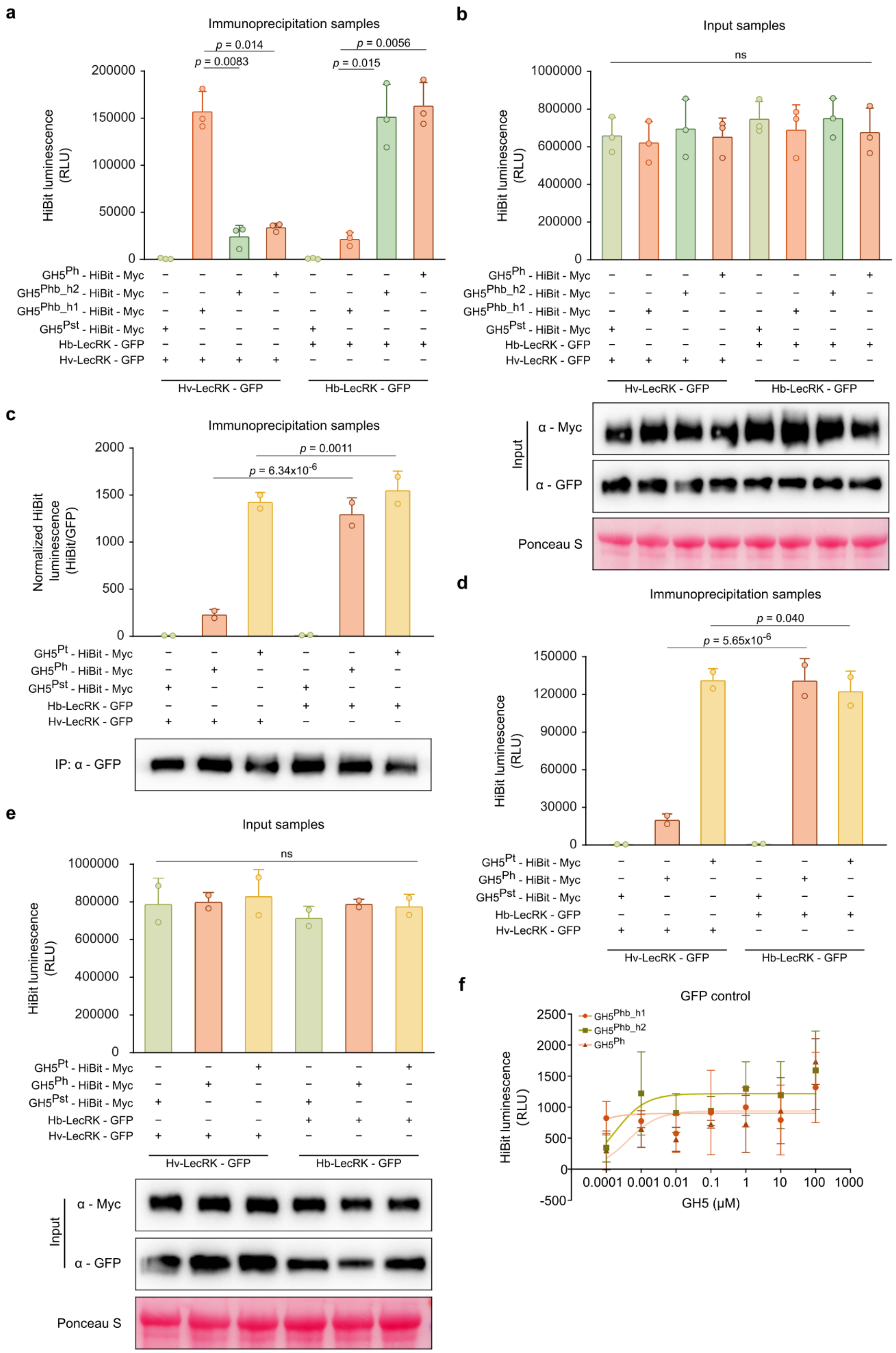
a
Immunoprecipitation samples
p = 0.014
p = 0.0083
p = 0.0056
p = 0.015
HiBit luminescence (RLU)
Hv-LecRK - GFP
Hb-LecRK - GFP
b
Input samples
ns
c
Immunoprecipitation samples
p = 0.0011
p = 6.34x10-6
Normalized HiBit luminescence (HiBit/GFP)
IP: α - GFP
d
Immunoprecipitation samples
p = 0.040
p = 5.65x10-6
e
Input samples
ns
Input
α - Myc
α - GFP
Ponceau S
f
GFP control
GH5Phb_h1
GH5Phb_h2
GH5Ph
GH5 (μM)

**Extended Data Fig. 6. HiBiT-based quantitative co-immunoprecipitation assay of GH5$^{Ph}$, GH5$^{Phb_h2}$, GH5$^{Phb_h1}$ GH5$^{Pst}$ and GH5$^{Pt}$ with Hv-LecRK or Hb-LecRK. a,b**, HiBiT-based quantitative co-immunoprecipitation assay of Myc-HiBiT-tagged GH5$^{Ph}$, GH5$^{Phb_h2}$, GH5$^{Phb_h1}$ and GH5$^{Pst}$ (as negative control) transiently co-expressed in *N. benthamiana* leaves with GFP-tagged full-length Hv-LecRK or Hb-LecRK. Absolute HiBiT luminescence values after background subtraction are shown for immunoprecipitated (IP) samples (**a**) and input samples (**b**, top). Protein abundance in input samples was detected by immunoblotting with anti-Myc and anti-GFP antibodies (**b**, bottom). Data are presented as mean ± s.d. normalized luminescence intensities from three independent experiments (n = 3). Circles indicate individual data points. The statistical significance of differences was determined by paired two-sample Student's *t*-test (two-tailed); *P*-values are indicated above the comparisons. Cases with no significant difference are indicated with “ns”. **c–e**, HiBiT-based quantitative co-immunoprecipitation assay of Myc-HiBiT-tagged GH5$^{Pt}$, GH5$^{Ph}$, and GH5$^{Pst}$ (as negative control) transiently co-expressed in *N. benthamiana* leaves with GFP-tagged full-length Hv-LecRK or Hb-LecRK. Total protein extracts were subjected to immunoprecipitation (IP) using anti-GFP beads. HiBiT luminescence (**c**, top) was measured from IP samples and normalized to the abundance of immunoprecipitated Hv-LecRK or Hb-LecRK, quantified by ImageJ analysis of anti-GFP immunoblots of the IP samples (**c**, bottom). Absolute HiBiT luminescence values after background subtraction are shown for IP samples (**d**), and input samples (**e**, top), with corresponding protein abundance in input samples detected by immunoblotting with anti-Myc and anti-GFP antibodies (**e**, bottom). Data are presented as mean ± s.d. normalized luminescence intensities from six technical replicates pooled from two independent experiments (n = 2). Circles indicate individual data points. The statistical significance of differences was determined by paired two-sample Student's *t*-test (two-tailed); *P*-values are indicated above the comparisons. Cases with no significant difference are indicated with “ns”. **f**, Negative control binding assay showing GFP incubated with increasing concentrations of HiBiT-tagged GH5$^{Ph}$, GH5$^{Phb_h1}$ and GH5$^{Phb_h2}$ proteins. The HiBiT luminescence was measured after incubation of GFP proteins with the increasing concentrations of GH5 proteins. Data represent mean ± s.e.m. from three independent experiments, with four technical replicates each.

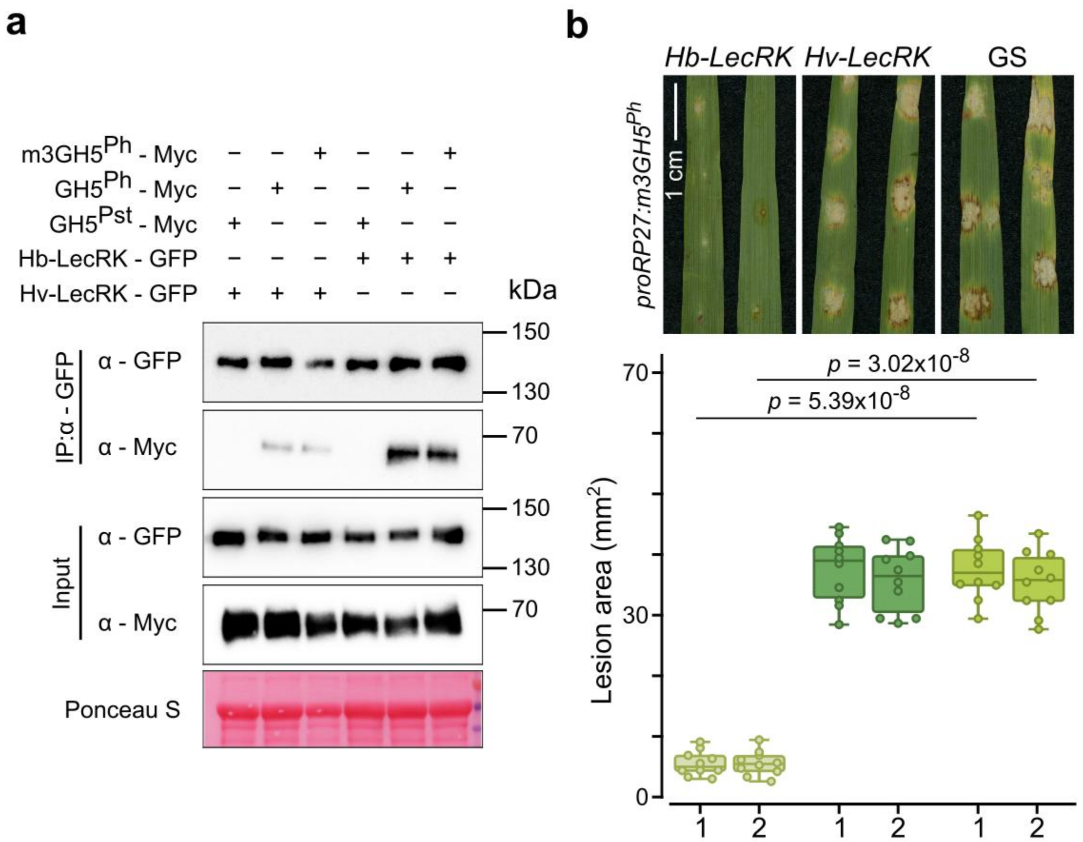


**Extended Data Fig. 7. Loss of catalytic activity does not alter GH5$^{Ph}$ specificity for LecRK. a**, *In planta* co-immunoprecipitation assay of Myc-tagged m3GH5$^{Ph}$ (GH5$^{Ph}$ variant E264A and E368A), GH5$^{Ph}$, and GH5$^{Pst}$ (as negative control) transiently co-expressed in *N. benthamiana* leaves with GFP-tagged full-length Hv-LecRK or Hb-LecRK. Total protein extracts were immunoprecipitated with anti-GFP beads (IP) and probed with anti-Myc and anti-GFP antibodies. **b**, Response of barley transgenic lines expressing *Hb-LecRK* or *Hv-LecRK*, or the wild type Golden SusPtrit (GS), after inoculation with *M. oryzae* expressing m3*GH5$^{Ph}$* driven by the *RP27* promoter. Representative leaf images (top) and quantification of lesion areas (bottom) at 6 days post-inoculation (dpi) are shown. Leaves were inoculated using the spot inoculation method. Numbers 1 and 2 on the *x*-axis represent two independent fungal transformants. Spot inoculations were performed on intact barley leaves without pricking, and photographs were taken at 6 dpi. Boxplots represent the mean lesion area ($n$ = 20) obtained from three leaves collected from three independent biological replicates for each treatment and experiment. Circles denote individual data points. The horizontal line denotes the median, box edges mark the first and third quartiles, and whiskers extend to the minimum and maximum values. The statistical significance of differences was determined by paired two-sample Student's $t$-test (two-tailed); $P$-values are indicated above the comparisons. ns, not significantly different. All experiments were repeated three times with consistent results.

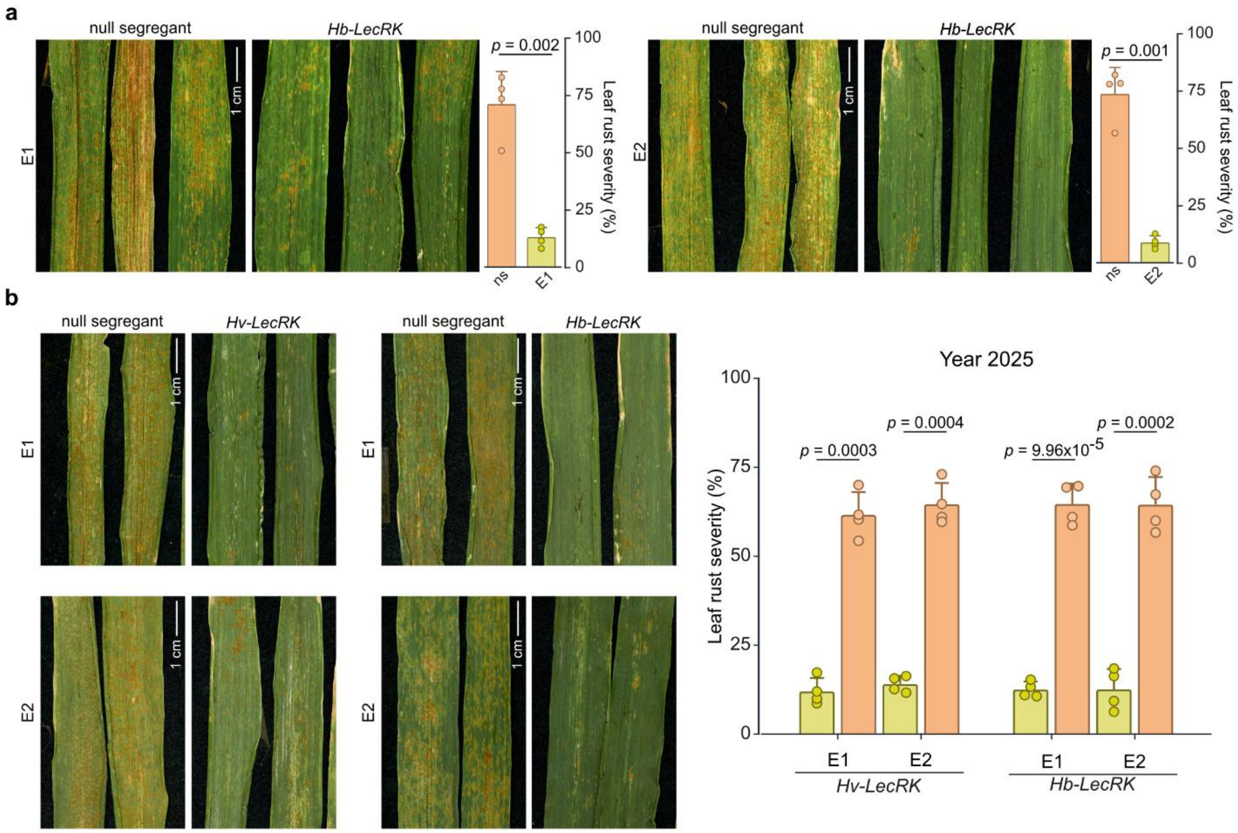


**Extended Data Fig. 8. Field evaluation of wheat transgenic lines expressing *Hv-LecRK or Hb-LecRK* for resistance to wheat leaf rust. a**, Representative photographs and quantification of leaf rust severity on the flag leaves from wheat transgenic lines expressing *Hb-LecRK* and corresponding null segregants (ns) in the 2024 field experiment. Disease severity was assessed 70 days after inoculation of the spreader rows with a mixture of seven *Pt* isolates. Values represent mean ± s.d. from four independent rows, each containing 15 plants (n = 60). E1 (left) and E2 (right) represent two independent transgenic events. **b**, Representative photographs (left) of leaf rust infection on the flag leaves from wheat transgenic lines expressing *Hv-LecRK* or *Hb-LecRK* and their corresponding null segregants from the 2025 field experiment. Quantification of leaf rust severity (right) on flag leaves of wheat transgenic lines expressing *Hv-LecRK* or *Hb-LecRK* and their corresponding null segregants in the 2025 field experiment. Disease severity was recorded 64 days after inoculation of spreader rows with seven mixed *Pt* isolates. Values represent mean ± s.d. from four independent rows, each containing 15 plants (n = 60). E1 and E2 represent two independent transgenic events. For all quantification data, statistical significance of differences was determined by paired two-sample Student's *t*-test (two-tailed).

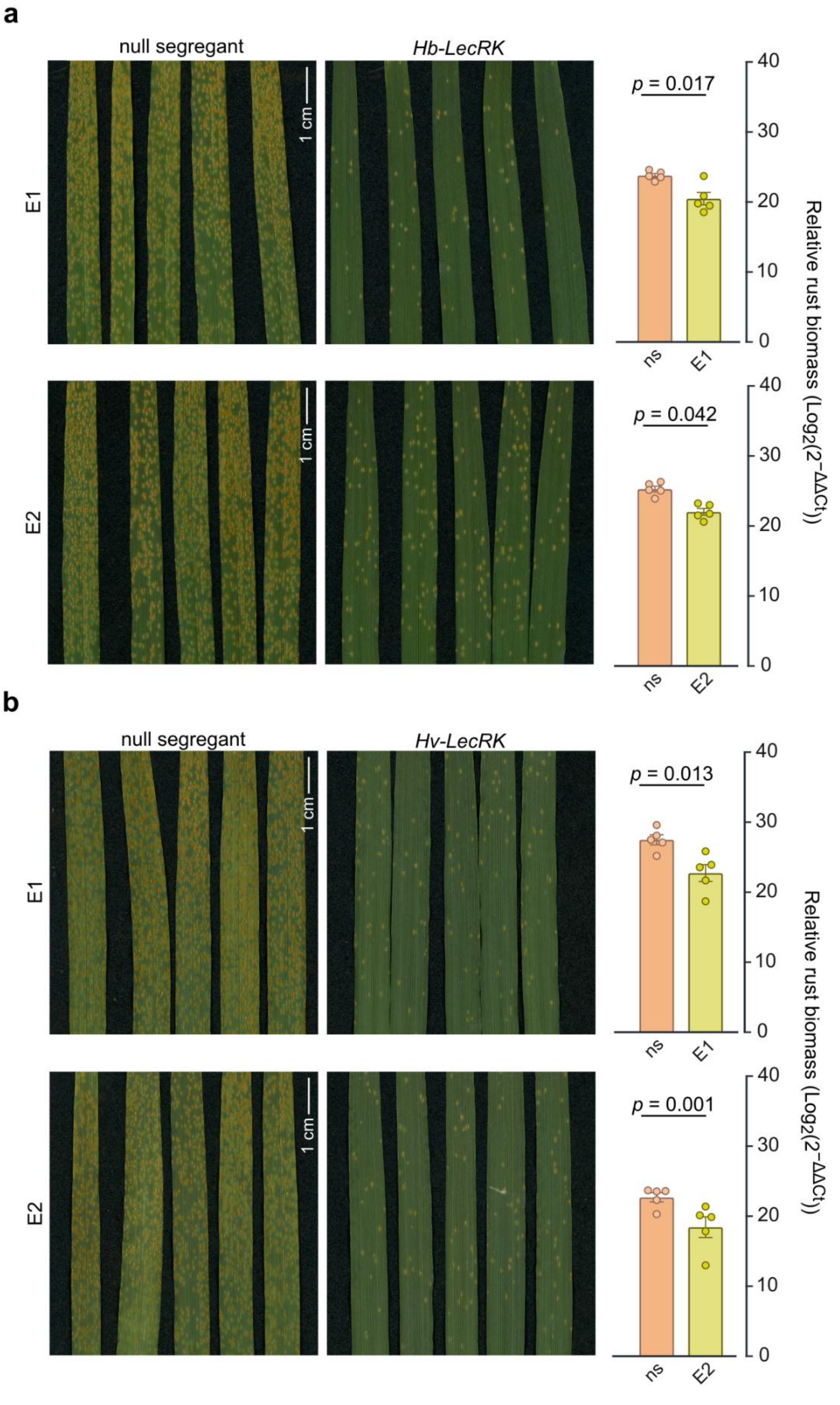

a
null segregant
Hb-LecRK
E1
E2
1 cm
p = 0.017
p = 0.042
ns
E1
E2
Relative rust biomass (Log2(2−ΔΔCt))
0
10
20
30
40
b
null segregant
Hv-LecRK
E1
E2
1 cm
p = 0.013
p = 0.001

**Extended Data Fig. 9. Wheat transgenic lines expressing *Hv-LecRK or Hb-LecRK* exhibit increased resistance to wheat leaf rust in controlled conditions. a**,**b**, Representative photographs (left) of leaves from adult wheat transgenic lines expressing *Hb-LecRK* (**a**, top) or *Hv-LecRK* (**b**, bottom) and their null segregants (ns) after inoculation with *Pt* isolate B9414. The photographs were taken at 14 dpi for five independent biological replicates ($n$ = 5) and two independent events (E1 and E2). Relative fungal biomass (right) quantified in *Hb-LecRK* (**a**, top) or *Hv-LecRK* (**b**, bottom) transgenic lines and null segregants using qPCR ($n$ = 5). Values are shown as means ± standard deviation. The statistical significance of differences was determined by paired two-sample Student's $t$-test (two-tailed). These experiments were performed under controlled laboratory (growth chamber) conditions. The experiments were performed three times with consistent results each time.

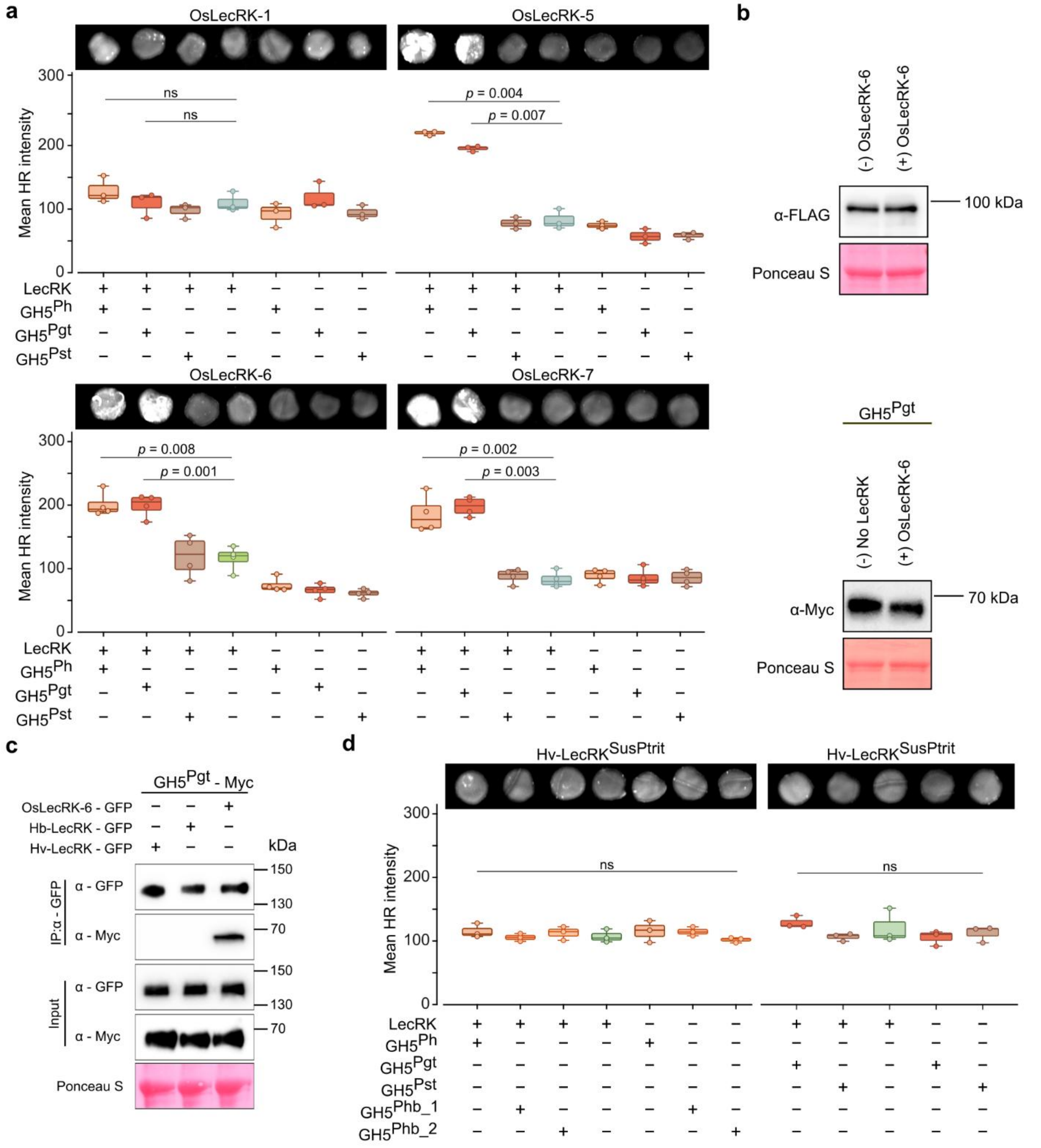


**Extended Data Fig. 10. HR response after co-expression of *OsLecRKs* and *Hv-LecRK*$^{SusPtrit}$ with *GH5s* from different rust pathogens. a**, HR imaging (top) and quantification (bottom) in *N. benthamiana* leaves co-expressing *GH5*$^{Ph}$, *GH5*$^{Pgt}$ and *GH5*$^{Pst}$ with *OsLecRK-1*, *OsLecRK-5, OsLecRK-6,* or *OsLecRK-7*. Corresponding *GH5s* and *OsLecRKs* expressed alone are shown as controls. HR was imaged at 3 dpi (Fusion-FX imager). Box plots represent the quantification of mean HR intensity (light orange, GH5$^{Ph}$; dark brown, GH5$^{Pst}$; salmon, GH5$^{Pgt}$; expressed with *OsLecRKs* or alone) from four biological replicates ($n = 4$). Circles indicate individual data points. The horizontal line denotes the median, box edges mark the first and third quartiles, and whiskers extend to the minimum and maximum values. The statistical significance of differences was determined by paired two-sample Student's *t*-test (two-tailed); *P*-values are indicated above the comparisons. Cases with no significant difference are indicated with “ns”. **b**, Immunoblots showing protein abundance of OsLecRK-6 and $GH5^{Pgt}$. Total protein was extracted at 36 h post-infiltration and expression of OsLecRK-6 was detected by anti-FLAG antibodies and the expression of $GH5^{Pgt}$ was detected by anti-Myc antibodies. (+) indicates co-expression of lectin receptor kinase with GH5, (-) indicates infiltration of lectin receptor kinase alone, ‘No LecRK’ indicates GH5 expressed alone. **c**, *In planta* co-immunoprecipitation assay of Myc-tagged $GH5^{Pgt}$ transiently co-expressed in *N. benthamiana* leaves with GFP-tagged full-length OsLecRK-6 or Hb-LecRK or Hv-LecRK. Total protein extracts were immunoprecipitated with anti-GFP beads (IP) and probed with anti-Myc and anti-GFP antibodies. **d**, HR imaging (top) and quantification (bottom) in *N. benthamiana* leaves co-expressing $GH5^{Ph}$, $GH5^{Pgt}$ and $GH5^{Pst}$ with $Hv\text{-}LecRK^{SusPtrit}$. Corresponding *GH5s* and *LecRK* expressed alone are shown as controls. HR was imaged at 3 dpi (Fusion-FX imager). Box plots represent the quantification of mean HR intensity from three biological replicates ($n$ = 3). Circles indicate individual data points. The horizontal line denotes the median, box edges mark the first and third quartiles, and whiskers extend to the minimum and maximum values. The statistical significance of differences was determined by paired two-sample Student's *t*-test (two-tailed). Cases with no significant difference are indicated with “ns”. The experiments were performed three times with consistent results each time.
