## Supplementary Note 1 for "Host specificity in cereal rust fungi is mediated by a conserved glycoside hydrolase family"

***Plant material.*** 'Vada' is a West European barley cultivar, previously released by the Department of Plant Breeding, Wageningen Agricultural University, and has a high level of partial resistance to *P. hordei*[1].

***DNA extraction for long-read sequencing.*** High molecular weight DNA was extracted from dark-treated leaves of Vada at the seedling stage and sent to the Arizona Genomics Institute and the French Plant Genomic Resource Center (INRAE-CNRGV) for HMW genomic DNA extraction, PacBio HiFi library preparation, and sequencing of 4 SMRT cells from each sequencing platform. Briefly, DNA purity was assessed on a NanoDrop NP-1000 spectrophotometer (NanoDrop Technologies), DNA concentration was measured with a Qubit dsDNA high-sensitivity assay and DNA size was validated by pulsed-field gel electrophoresis (PFGE). Sequencing libraries were constructed following the manufacturer's protocol using a SMRTbell Express Template Prep Kit 2.0. Libraries were quantified using the Qubit dsDNA high-sensitivity assay, and size was checked on a Femto Pulse System (Agilent). Sequencing was performed on PacBio Sequel II systems in CCS mode for 30 h.

***RNA-seq library preparation and sequencing.*** Around 100 mg of frozen and ground tissue from roots, whole aerial parts at seedling stage, flag leaves, fully emerged spikes, glumes, and grains was used for RNA isolation with a Maxwell RSC Plant RNA Kit (AS1500) with a Maxwell RSC48 instrument following the kit protocol (Promega). For RNA-seq, around 10 Gb of Illumina 150-bp paired-end reads were generated for each tissue.

***Optical map production.*** The same dark-treated Vada leaves collected at seedling stage were used for the PacBio sequencing and Ultra-HWM DNA isolation using the Plant DNA Isolation Kit protocol (Bionano Genomics) at the French Plant Genomic Resource Center (INRAE-CNRGV). Labelling was performed using direct labelling enzyme (DLE1) and staining of the HMM DNA according to the Bionano Prep Direct Label and Stain (DLS) protocol (30206-Bionano Genomics). Optical maps were generated using the Bionano Genomics Saphyr System (Saphyr Chip G1.2) according to the Saphyr System User Guide (3024-Bionano Genomics). Data processing was performed using the Bionano Solve v.3.6 software (https://bionanogenomics.com/support/software-downloads).

***Omni-C library.*** The Omni-C library was prepared and sequenced from the dark treated young leaves of Vada at the National Genomics Infrastructure (NGI)/Uppsala Genome Center and using the Dovetail Omni-C Kit according to the manufacturer's protocol. In brief, chromatin was fixed in place in the nucleus. Fixed chromatin was digested with DNase I and then extracted. Chromatin ends were repaired and ligated to a biotinylated bridge adapter, before proximity ligation of adapter containing ends. After proximity ligation, crosslinks were reversed, and the DNA was purified from proteins. Purified DNA was treated to remove biotin that was not internal to ligated fragments. Four sequencing libraries were generated using Illumina-compatible adapters. Biotin-containing fragments were isolated using streptavidin beads before PCR enrichment of the library. One library was sequenced on an Illumina NovaSeq 6000 S4 platform to generate 1,292 million $2 \times 150$ bp read pairs.

***Genome assembly and scaffolding.*** The PacBio HiFi reads (38-fold coverage) were assembled using hifiasm (v.0.16.1)[2] with default parameters. Hybrid scaffolding incorporating the PacBio contigs and the optical map (~124-fold coverage, molecule N50 of 224 kb) was performed using the hybridScaffold pipeline (Bionano Solve 3.6) with default parameters. The pseudomolecules were assembled using the Omni-C reads processed with Juicer tools (https://github.com/aidenlab/juicer - v.1.6) and 3D-DNA (https://github.com/aidenlab/3d-dna - v.180114). In brief, the preprocessing of the Omni-C reads was performed with juicer.sh (parameter: -s none). Juicebox (https://github.com/aidenlab/Juicebox - v.1.11.08) was used to visualize the Hi-C map and to perform the manual curation for final barley cv. Vada assembly. To validate the genome assembly, we remapped the optical map onto the pseudomolecule using the hybridScaffold pipeline (Bionano Solve 3.6).

This final assembly of 4.3 Gb comprised the seven chromosomes (4.2 Gb) and one unanchored chromosome (156.7 Mb) containing the unassigned contigs and scaffolds. An alignment of Vada pseudomolecules against Barley cv Morex V3 with CHROMEISTER (v1.5.a) revealed a high collinearity for each chromosome and the BUSCO (Benchmarking Universal Single-Copy Orthologs) analysis indicated a high completeness with a score of 97.8%.

***Annotation.*** Gene model prediction was performed following the method described by Athiyannan, *et al.*[3] with minor modifications, combining transcriptomics data, *ab initio* prediction and protein homology. First, RNA-seq data from the six tissues were mapped to their respective reference assemblies using STAR[4] (v2.7.0f; parameters: --outFilterMismatchNoverReadLmax 0.02) and assembled into transcripts with Stringtie[5] (v2.1.4; parameters: --rf -m 150 -f 0.3 -t ). The transcripts from the six tissues were merged using Stringtie[5] (v2.1.4; parameters: --merge -m 150) into a pool of candidate transcripts, and Transdecoder (v5.5.0; https://github.com/TransDecoder/TransDecoder) was used to find potential open reading frames and to predict protein sequences within the candidate transcript set. For the *ab initio* gene predictions, we used BRAKER2 (v2.1.2)[6] and FgeneSH (v8.0.0; http://www.softberry.com). Briefly, BRAKER2 gene prediction was trained supported by RNA-seq (parameters: --softmasking --gff3 --cores=48 --nocleanup --bam='list of BAM files'). For the FgeneSH prediction, pseudomolecules were repeat masked using a *de novo* repeat library constructed with the EDTA pipeline[7] and the TREP database[8]. FgeneSH annotation was performed with the monocot matrix for gene prediction. For the protein homology evidences, we used the translated proteins from gene annotations of 19 accessions of the barley pan-genome[9] and Morex v3[10], the related grass species *Brachypodium distachyon*[11] and rice[12], and the *Triticeae* and *Poaceae* protein sequences downloaded from the UniProt database (2021_03). All protein sequences were mapped against the Vada assembly using GenomeThreader[13] (v1.7.1; parameters: -startcodon -finalstopcodon -species rice -gcmincoverage 70 -prseedlength 7 -prhdist 4 -gff3out). We used EVidenceModeler[14] (v1.1.1) to join all the gene evidences from transcriptomics, *ab initio* predictions, and protein alignments with weights adjusted according to the input source (FgeneSH= 2; BRAKER2 = 1; protein homology = 6; transcriptomics = 12). Finally, we performed two rounds of isoform and UTR prediction using the PASA pipeline (v2.5.1) with default parameters. Gene models were classified into high- and low-confidence according to classification criteria used by the International Wheat Genome Sequencing Consortium[15] and by Mascher, *et al.*[10]. Briefly, protein-encoding gene models were considered as complete when start and stop codons were present. A comparison against PTREP[8], UniPoa (Poaceae database of annotated proteins from UniProt_2021_03) and UniViri (Viridiplantae database) was performed using DIAMOND[16] (v2.0.9) and a BUSCO (v5.2.2) analysis against the poales database (v10; parameters: -m prot -c 20 -l poales). Gene candidates were further classified using the following criteria: A high confidence (HC) gene model is complete with a hit in the UniViri database and/or in UniPoa and/or BUSCO poales database but not in PTREP. A low confidence (LC) gene model is incomplete and has a hit in the UniViri or UniPoa or BUSCO poales database but not in PTREP, or the protein sequence is complete with no hit in UniViri / UniPoa / BUSCO poales / PTREP. Putative functional annotations were assigned to HC and LC transcripts using a protein comparison against the UniProt database (2021_03) and PFAM domain signatures, and GO were assigned using InterproScan version 5.55-88.0[17]. The final annotation for barley cv. Vada consists of 39,075 HC genes and 96,805 LC genes (BUSCO scores of 99.1% for HC genes).

The authors would like to acknowledge support of the National Genomics Infrastructure (NGI)/Uppsala Genome Center (https://www.uu.se/en/research/research-infrastructure/national-research-infrastructures-in-which-uppsala-university-participates/ngi) for providing assistance in NGS sequencing and of the Genotoul bioinformatics platform Toulouse Midi-Pyrenees (Bioinfo Genotoul, doi: 10.15454/1.5572369328961167E12, http://bioinfo.genotoul.fr) for providing computing resources. We thank AGI and INRAE-CNRGV for sequencing support.
